# Glyoxal induces DNA-Protein Crosslinking in Cells

**DOI:** 10.64898/2026.07.30.741824

**Authors:** Krishna C. Gurajala, Elijah M. Barnes, Luke Erber

## Abstract

Glyoxal (GO) is a small, highly reactive molecule that is produced naturally in cells during normal metabolism and can also come from processed foods and oxidative stress. Because of its high reactivity, glyoxal can modify DNA and proteins to form harmful products called advanced glycation end-products (AGEs), which have been linked to diseases such as diabetes, cancer, and aging. Although glyoxal is known to modify DNA and proteins, it is not well understood whether it can form DNA-protein crosslinks (DPCs), a type of DNA damage in which proteins become permanently attached to DNA. In this study, we investigated glyoxal induced DPC formation in HeLa cells using biochemical assays and mass spectrometry-based proteomics experiments. We observed that glyoxal exposure elevated cellular DPC formation in a concentration- and time-dependent manner. Cells with reduced SPRTN expression accumulated higher levels of DPCs, suggesting that SPRTN plays an important role in repairing glyoxal induced DNA damage. Proteomics experiments revealed 469 proteins exhibited elevated DNA association in glyoxal-treated samples, including histones and other proteins involved in chromatin organization, DNA replication, DNA repair, and gene expression. *In-vitro* experiments confirmed that glyoxal can directly crosslink DNA with histone proteins. Overall, this study provides the first evidence that glyoxal forms DNA-protein crosslinks in human cells. These findings provide a foundation for future studies on the chemical structure, biological effects and repair of glyoxal induced DNA-protein crosslinks and their possible role in human disease.

## INTRODUCTION

DNA-protein crosslinks (DPCs) are bulky lesions which form when proteins become covalently attached to DNA. These lesions interfere with many essential cellular processes, including DNA replication, transcription, recombination, chromatin remodeling, and DNA repair (1). Because the protein remains physically attached to DNA, DPCs create a major obstacle for enzymes that need access to genetic material. If these lesions are not efficiently removed, they can cause replication stress, genome instability, mutations, and cell death. Cells therefore rely on specialized repair pathways, including the metalloprotease SPRTN and the proteasome, to degrade the protein component before the remaining DNA lesion can be repaired (2-4). Polymorphisms in the SPRTN gene defines a genetic disorder, Ruijs-Aalfs syndrome. Individuals with Ruijs-Aalfs syndrome exhibit elevated DPC levels, genomic instability, accelerated aging, and greater risk for hepatocellular carcinoma (4-6).

DPCs are formed by exposure to endogenous and exogenous agents. Environmental chemicals (7), ultraviolet light (8), transition metals (9), and several anticancer drugs such as cisplatin, adriamycin, and camptothecin (10-12) are known to generate DPCs. In addition to these external sources, DPCs are produced by reactive metabolites generated by cellular metabolism (13, 14). These endogenous lesions are of particular interest because they accumulate throughout life and may contribute to aging and cancer (3).

Among endogenous metabolites, glyoxal (GO) is a simple reactive dicarbonyl compound produced in cells. Glyoxal is generated during glucose autooxidation (15) and lipid peroxidation (16). Additional exposure occurs through processed foods, cigarette smoke, cosmetics and disinfectants (17, 18). Under normal physiological conditions, glyoxal is present at low concentrations with free glyoxal levels reported in the nanomolar range in plasma (∼100 -120 nM) and approximately ∼0.1-1 µM inside the cell (19). The earliest studies using derivatization-based methods reported much higher total glyoxal levels in plasma (∼215 µM in healthy individuals and up to ∼469 µM in poorly controlled diabetes), although these values are likely overestimated (20). These relatively low levels of glyoxal are maintained primarily through the glutathione-dependent glyoxalase pathway, in which glyoxalase I (GLO1) and glyoxalase II (GLO2) convert glyoxal into less reactive metabolites such as glycolate **(Figure 1)** (21). Consistent with this role, loss of GLO1 expression has been associated with increased accumulation of glyoxal-derived adducts, including carboxymethyllysine (CML) and carboxymethylarginine (CMA) (22), while GLO2 deficiency has been linked to reduced glycolate formation (23). In addition to the glyoxalase system, alternative detoxification pathways involving aldehyde dehydrogenases and aldose/aldehyde reductases have also been reported (21).

**Figure 1:**
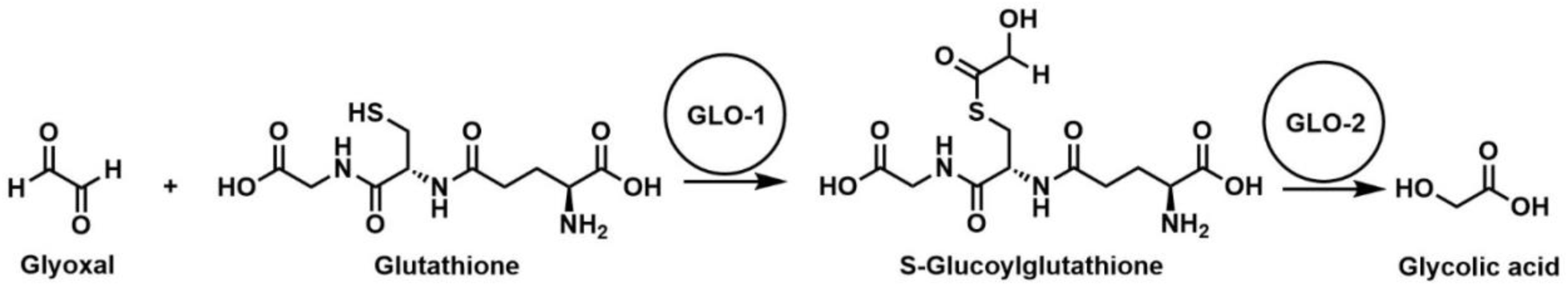
Detoxification pathway of glyoxal by glyoxalase I (GLO1) and glyoxalase II (GLO2).

The high reactivity of glyoxal results in modification of many different biomolecules. In proteins, glyoxal mainly reacts with lysine, cysteine and arginine residues, forming stable glycation products, including Nε-(carboxymethyl) lysine (CML), S-carboxymethylcysteine (CMC), and Nε- (carboxymethyl)arginine (CMA), as well as hydroimidazolone derivatives (G-H1, G-H2, and G-H3) (**Figure 2**). In DNA, glyoxal reacts mainly with guanine bases to form adducts such as dG-GO and CMdG (24). Furthermore, due to its bifunctional aldehyde structure, glyoxal is not limited to forming simple adducts but can also bridge two nucleophilic sites, leading to crosslink formation. In proteins, glyoxal mediates protein-protein crosslinks through reactions between lysine and arginine residues, generating well-characterized advanced glycation end products such as glyoxal lysine dimer (GOLD), glyoxal-derived imidazolium crosslink (GODIC), and glyoxal lysine amidine (GLA) (25, 26). Similarly, in nucleic acids, glyoxal can induce DNA-DNA crosslinks, including dG-gx-dC and dG-gx-dA adducts, by reacting with guanine bases on opposing strands (27). These crosslinking reactions further highlight the ability of glyoxal to simultaneously interact with two nucleophiles, contributing to structural damage, genomic instability, and impaired cellular function.

**Figure 2:**
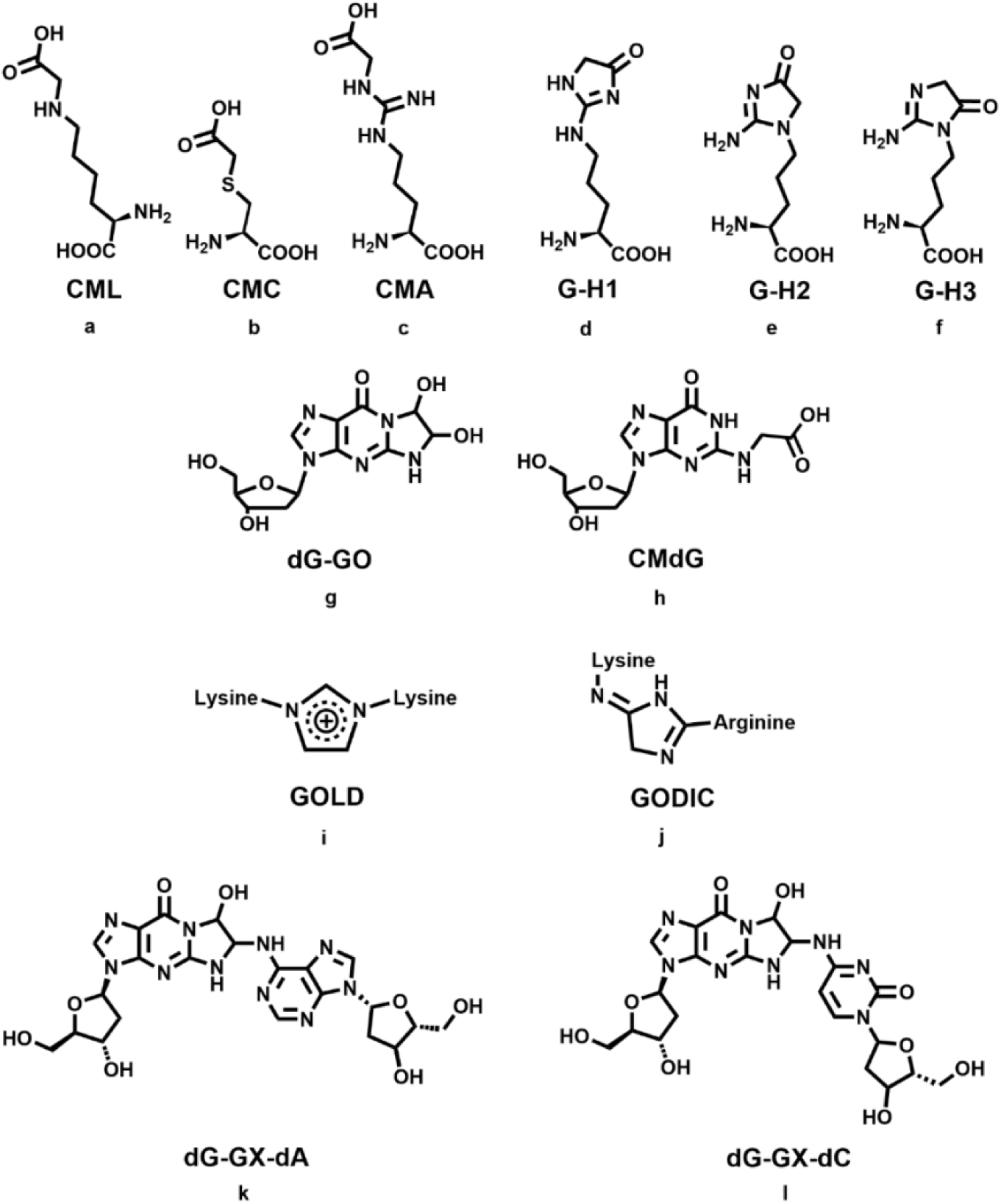
Structures of glyoxal-derived protein and DNA adducts. a) Nε-(carboxymethyl)lysine (CML), b) S-(carboxymethyl)cysteine (CMC), c) Nε- (carboxymethyl)arginine (CMA), d-f) Hydroimidazolone adducts formed from glyoxal: G-H1, G-H2, G-H3, g) deoxyguanosine-Glyoxal adduct (dG-GO), h) N2-carboxymethyl-2′-deoxyguanosine (CMdG), i) glyoxal lysine dimer (GOLD), j) glyoxal-derived imidazolium crosslink (GODIC), k-l) DNA-DNA crosslinks formed via glyoxal: dG-gx-dA and dG-gx-dC.

Although glyoxal-derived protein adducts, protein-protein crosslinks, DNA adducts, and DNA-DNA crosslinks have been extensively characterized, much less is known about glyoxal-induced DNA-protein crosslinks. An early in-vitro study demonstrated that reactive dicarbonyl compounds such as glyoxal and methylglyoxal can induce DNA-protein crosslinks (DPCs). Using purified DNA polymerase and DNA substrates, this study showed that both glyoxal and methylglyoxal formed covalent DNA-protein complexes, although methylglyoxal exhibited greater crosslinking efficiency than glyoxal (13). While this finding established the chemical feasibility of dicarbonyl-induced DPC formation, this was limited to a single model protein and did not address whether glyoxal generates DPCs with diverse DNA-binding proteins in living cells. Consequently, the extent of glyoxal-induced DPC formation, the identities of proteins involved, and the biological pathways affected remain largely unknown.

In this study, we investigated DNA-protein crosslink formation following glyoxal exposure in human cells. We combined complementary biochemical assays, including the advanced recovery of K-SDS precipitates (ARK) assay and the superior method for true DNA-protein crosslink recovery (STAR) assay, to quantify DPC formation and isolate DNA crosslinked proteins. Mass spectrometry-based proteomics and bioinformatic analyses were applied to identify proteins enriched within glyoxal-induced DPCs and determine the biological pathways affected by glyoxal exposure. Together, our results demonstrate that glyoxal is an inducer of DNA-protein crosslinks and provide new insight into the proteins and cellular processes targeted by glyoxal-derived DNA damage.

## EXPERIMENTAL SECTION

### Reagents and Materials

Glyoxal (40 wt. % in H2O; Cat. No. A716938), guanidine hydrochloride (Cat. No. A741759), and 1,2-cyclohexanedione (Cat. No. A199204) were purchased from Ambeed. Recombinant human Histone H2A (Cat. No. 31490), Histone H2B (Cat. No. 31492), Histone H3.1 (Cat. No. 31294) and Histone H2A antibody (Cat. No. 39236) were obtained from Active Motif.

Dulbecco’s Modified Eagle Medium (DMEM; Cat. No. 11320-033), Trypsin-EDTA (0.25%; Cat. No. 25200056), sodium dodecyl sulfate (SDS; Cat. No. 15525017), PrestoBlue™ Cell Viability Reagent (Cat. No. P50200), Quant-iT™ PicoGreen™ dsDNA Reagent (Cat. No. P7581), Pierce™ Silver Stain Kit (Cat. No. 24612), crystal violet (Cat. No. B21932.14), and NuPAGE 4–12% BisTris SDS-PAGE gels (Cat. No. NP0321BOX) were purchased from Thermo Fisher Scientific. Fetal bovine serum (FBS; Cat. No. A1620) was purchased from Biowest.

Proteinase K (Cat. No. P8107S) was obtained from New England Biolabs. Sodium chloride (Cat. No. S9888), guanidine isothiocyanate (Cat. No. 5120-OP), Sarkosyl (Cat. No. 61747), Triton X100 (Cat. No. X100), potassium chloride (Cat. No. P3911), iodoacetamide (Cat. No. I1149), and 2,5-dioxopyrrolidin-1-yl acetate (Cat No. SY3H3249F55C-1G) were purchased from Sigma Aldrich.

Ethylenediaminetetraacetic acid (EDTA; Cat. No. E478-500), dithiothreitol (DTT; Cat. No. D10711G), and ethanol (Cat. No. 07-678-007) were purchased from Fisher Scientific. Sodium deoxycholate (Cat. No. 97062-024) was purchased from VWR.

### Cell Lines

Wildtype and SPRTN-knockdown HeLa cells were previously prepared in the Erber lab(28). GLO1 knockout HEK293 cells were received from Dr. James Galligan’s laboratory(29). These cell lines were maintained in Dulbecco’s modified Eagle medium (DMEM) supplemented with 10% FBS and 1% penicillin/streptomycin antibiotics. All cells were maintained in a humidified incubator at 37 °C with 5% CO2 and grown until a confluency of ∼80% or higher was achieved. Cell culture experiments were executed within a third to eighth passage range.

### Presto Blue Cell Viability Assay for Glyoxal Toxicity

Presto blue assay (30) was performed on HeLa cells according to the manufacturer’s instructions. The cells were seeded 10,000 per well into a 96-well plate for 24 hours, the media was replaced with glyoxal containing serum-free media for either 2 hours or 6 hours. After the treatment, glyoxal containing media was replaced with 100 µL of 10% presto blue reagent dissolved in serum-free DMEM. The 96-well plate was further incubated for another 10 minutes at 37°C, and the fluorescence was measured at Ex/Em 560/590 nm using BMG CLARIOstar® Plus Multi-Mode Microplate Reader. The fluorescence signal for each treatment condition was normalized to the vehicle-treated control to calculate cell viability.

### *In-vitro* Characterization of DPCs Generated with Recombinant Histone Proteins

To evaluate crosslink formation, recombinant histone was incubated with a 3’ 6-carboxyfluorescein (FAM)-labeled single stranded oligonucleotide containing the telomeric repeat sequence (TTAGGG)_3_, in the presence or absence of glyoxal (GO) at physiological pH. After incubation, the samples are denatured and separated by SDS-PAGE. The gels were then analyzed using fluorescence imaging and protein staining (14).

### Quantitation of DPCs Using the Ark Assay

DPC-containing DNA was isolated by using ARK assay (31). The cells were lysed by using 950 µL pre-warmed lysis buffer (5.6 M Guanidine thiocyanate, 10 mM Tris-HCl (pH 6.5), 20 mM EDTA, 4% Triton X-100, 1% Sarkosyl & 1% dithiothreitol). The lysates were sheared by passing through 1 mL pipette. The DNA was precipitated with 950 µL of prechilled ethanol and centrifuged at 20000 x g for 10 min. After the aspiration of supernatant, the pellet was washed with wash buffer A containing 20 mM Tris-HCl pH 6.5, 150 mM NaCl and 50% ethanol. The DNA/DPC pellet was dissolved in 0.5 mL of pre-warmed 1% SDS, 20 mM Tris-HCl (pH 7.5) and incubated at 42°C for 6 min. The samples were sheared by passing through 25-gauge needle 5 times and the proteins were precipitated by using 0.5 mL buffer consisting of 200 mM KCl and 20 mM Tris-HCl (pH 7.5). The samples were precipitated by incubating on ice for 6 min and centrifuged. The supernatant was collected for DNA measurement. The DPC pellet was washed in 1mL of 100 mM KCl, 20 mM Tris-HCl (pH 7.5) and incubated at 55 °C for 5 min, followed by incubation on ice and centrifuged at 20000 x g. The supernatant was collected and combined with the previous fraction. The wash procedure was repeated one more time and the pellet was dissolved in 1mL of proteinase K buffer containing 100 mM KCl, 20 mM Tris-HCl (pH 7.5) and 10 mM EDTA. The proteins were digested by adding proteinase K to a final concentration of 0.2 mg/mL and incubated at 37 °C overnight. After the digestion, the samples were centrifuged to remove any debris and the supernatant containing DPC associated DNA was collected to determine the DPC coefficient. The total DNA amounts were determined using the Quant-iT™ PicoGreen™ dsDNA reagent according to the manufacturer’s instructions. DPC levels were quantified by calculating the DPC coefficient, defined as the ratio of DNA recovered from the DPC-associated fraction to the sum of DNA recovered from the DPC-associated fraction and the total input DNA fractions. Results were expressed as fold change relative to untreated controls.

### DNA Quantitation Using Quant-it™ PicoGreen™ dsDNA Assay

We measured DNA in the input and DPC samples using the Quant-iT™ PicoGreen™ dsDNA reagent. In each well of a 96 well plate, we added a total of 100 µL, consisting of 90 µL of dye solution and 10 µL of sample or standard. The standard curve was prepared by diluting lambda DNA stock solution (0.3 µg/µL). After adding the samples and standards to the dye, the plate was incubated in the dark for 5 minutes. Fluorescence was measured at 480 and 520 nm wavelengths using a BMG CLARIOstar® Plus Multi-Mode Microplate Reader.

### STAR (Superior Method for True DNA-Protein Adducts Recovery) Assay

DNA-crosslinked proteins were isolated using STAR assay (32). In brief, HeLa cells (7 million), treated with 0, 0.1, 1, 5, 10 and 20 mM glyoxal for 2 hours. Cells were lysed with buffer A, consisting of 50 mM Tris-HCl (pH: 7.4), 1 mM EDTA, 150 mM NaCl, 1% Triton X-100, 0.5% deoxycholate (DOC), and 0.1% sodium dodecyl sulphate (SDS). Lysates were centrifuged to aspirate the supernatant. The insoluble pellet was solubilized in buffer B, consisting of 6 M guanidinium-HCl, 10 mM Tris-HCl (pH: 6.5), 20 mM EDTA, 4% Triton X-100, 0.1% SDS and 1% Dithiothreitol (DTT). DNA and associated DNA-protein crosslinks were isolated by ethanol precipitation. The precipitated material was washed three times with 75% ethanol to remove residual salts, detergents, and non-crosslinked proteins. The DNA containing pellet was resuspended in 16 mM NaOH and neutralized with 40 mM Tris-HCl (pH 7.0). DNA concentration was determined using the PicoGreen dsDNA quantitation assay (ThermoFisher) according to the manufacturer’s instructions. After measuring DNA concentration, samples were normalized to equal DNA concentrations prior to SDS-PAGE and western blot analysis.

### SDS-PAGE Separation and Total Protein Staining

To separate and visualize DNA-crosslinked proteins, equal amounts of DNA were heated at 90 °C in denaturing NuPAGE LDS buffer for 10 minutes and loaded onto 4-12% NuPAGE SDS-PAGE gels. DPC proteins were separated for 90 min at 120 V. Gels were washed with Milli-Q Water, stained by using the Pierce silver staining kit (ThermoFisher) according to the manufacturer’s instructions.

### Dot-Blot Procedure

DPC samples isolated using the STAR assay were added onto a nitrocellulose membrane using a Bio-Rad dot blot setup. The membrane was first pre-wetted with 1X TBS buffer, and samples were bound to the membrane by gravity. The membrane was washed with TBST and blocked with 3% milk in TBST for 1 hour. After blocking, the membranes were incubated with primary antibodies against H2A (rabbit, Active Motif, Cat. No. 39236) and dsDNA (mouse, HYB331-01), followed by appropriate secondary antibodies (Invitrogen’s goat anti-rabbit Alexa Fluor 647 and goat anti-mouse Alexa Fluor 680). The membranes were analyzed using an Odyssey Li-COR system.

### Proteomic Analysis of DNA-Protein Crosslinks DPC Isolation and Sample Preparation

DNA-Protein crosslinks (DPCs) were isolated by STAR assay from three control samples and three samples treated with 20 mM glyoxal. The samples were dried using speedvac and resuspended in 100 µL of 50 mM triethylammonium bicarbonate (TEAB) containing 2 mM CaCl_2_. Disulfide bonds were reduced by adding tris(2-carboxyethyl) phosphine (TCEP) to a final concentration of 5 mM and incubating at 55 °C for 30 minutes. Cysteine residues were then alkylated using iodoacetamide (IA) at a final concentration of 10 mM by incubating at room temperature in the dark for 30 minutes. Proteins were digested by adding 500 ng of trypsin, which specifically cleaves at lysine and arginine residues, by incubating overnight at 37 °C at 500 rpm. Peptide samples were purified using Pierce C18 spin columns, dried, and resuspended in 0.1% formic acid prior to LC-MS/MS analysis.

### LC-MS/MS Analysis

DPC peptide samples were analyzed using a Vanquish Neo nano-UPLC system coupled to an Orbitrap Ascend Tribrid mass spectrometer equipped with a FAIMS pro interface (Thermo Fisher Scientific). Peptides were separated on an Aurora Elite XT C18 analytical column (75 µm × 150 mm, 1.7 µm particle size; IonOpticks) using a reverse-phase gradient from 2-25% acetonitrile over 100 min followed by a 40% acetonitrile over 20 min. Data was acquired using data-dependent acquisition (DDA). Full MS scans were collected in the Orbitrap at 120,000 resolving power over an m/z range of 375-1500. Precursor ions with charge states of +2 to +7 were fragmented by higher-energy collision dissociation (HCD) using 28% normalized collision energy. FAIMS compensation voltages were cycled at -45 V, -60 V, and -75 V.

### Proteomics Data Processing and Statistical Analysis

Raw mass spectrometry data was processed using Proteome Discoverer 3.0 with SeQuest algorithm and searched against the human UniProt proteome database downloaded on March 21, 2025, together with a common contaminants database. Protein abundances were determined using label-free quantitation based on total peptide abundance. False discovery rates were controlled using Percolator with strict and relaxed FDR thresholds of 0.01 and 0.05, respectively. Proteomics data were further analyzed using Perseus software. Data were log-transformed and filtered to retain proteins with at least three valid values in at least one experimental group. Missing values were imputed using a normal distribution-based approach (downshift of 1.8 standard deviations from the mean with a width of 0.3). Correlation between biological replicates was assessed using multi-scatter plots and Pearson correlation analysis. Differential enrichment analysis between glyoxal-treated and control samples was performed using two-sample t-test with permutation-based false discovery rate (FDR) correction in Perseus (FDR = 0.05, S0 = 0.1, 250 randomizations). For volcano plot visualization in VolcaNoseR, proteins with log2fold change ≥ 2 or ≤ -2 and -log10(p-value) ≥ 1 were set as significantly changed proteins. The mass spectrometry proteomics data have been deposited to the ProteomeXchange Consortium via the PRIDE partner repository (33) with the dataset identifier PXD081900

### Statistical Analysis

Graphpad PRISM (version 10.1.0) was used to complete statistical analysis. Comparisons were completed with ordinary one-way ANOVA using Turkey’s multiple comparisons test, with a single pooled variance. Group datasets statistics were assessed using Ordinary two-way ANOVA using uncorrected Fisher’s LSD, with a single pooled variance. GP: 0.1234 (ns), 0.0332 (*), 0.0021 (**), 0.0002 (***), <0.0001 (***).

## RESULTS

### Glyoxal induces DNA-protein crosslinks in human cells

To determine whether glyoxal induces DNA-protein crosslinks (DPCs) in human cells, we first established treatment conditions that produced measurable DPC formation while maintaining cell viability. HeLa cells were exposed to increasing concentrations of glyoxal for 2 hours **(Figure 3A)** or 6 hours **(Figure S1)**, and cell viability was determined using PrestoBlue assay. Cell viability decreased in a concentration and time-dependent manner, with EC50 values of approximately 29.9 mM (2 hours) and 6.9 mM (6 hours). Based on these results, a maximum of 20 mM glyoxal for 2 hours was selected for subsequent experiments to limit excessive cell death. Similar studies evaluating glyoxal-induced DNA and protein modifications reported using low micromolar levels with multiday exposure conditions and millimolar levels with 8+ hours treatment conditions(34, 35). Millimolar levels of glyoxal are used to mimic acute intoxication and cytotoxicity(19).

**Figure 3:**
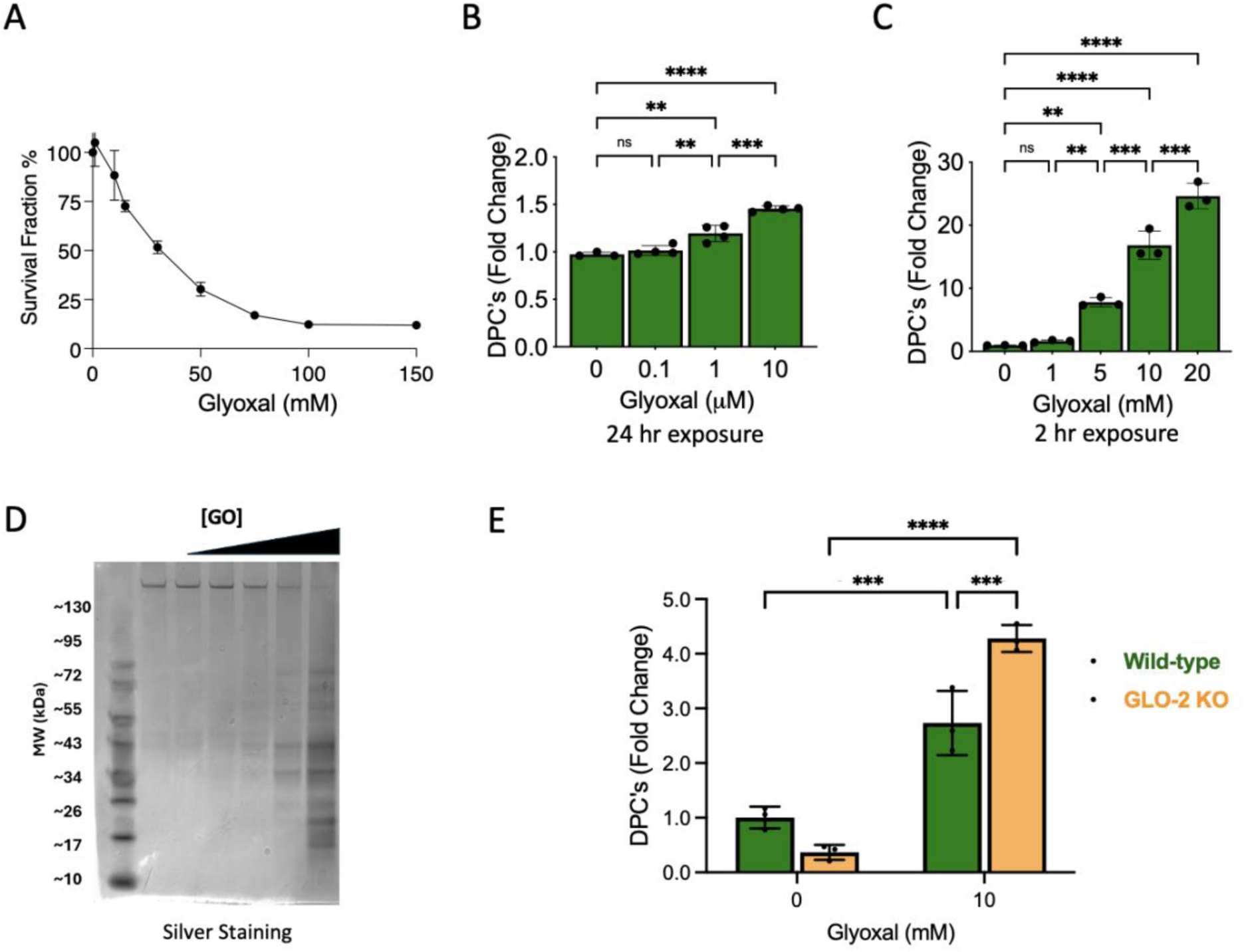
Glyoxal induces DNA-protein crosslinks in human cells. A) HeLa cells were treated with different concentrations of glyoxal (0,1,10, 15, 30, 50, 75, 100, and 150 mM) for 2 h and cell viability was measured using the PrestoBlue assay. Data represents the mean ± SD of three independent replicates (n=3). Nonlinear regression analysis was performed using GraphPad Prism. B) DPC levels (shown as fold change) were measured using the ARK assay following glyoxal exposure to micromolar concentrations for 24 hours. and C) millimolar concentrations for 2 hours D) HeLa cells were treated with 0, 0.1, 1, 5, 10, or 20 mM GO for 2 hours and were processed using STAR assay to isolate DPCs. Isolated DPCs were resolved by 4-12% SDS-PAGE and visualized by the silver staining. E) DPC levels (shown as fold change) were measured in wild type HEK293 and GLO-2 KO cells treated with glyoxal (10 mM, 2 hours). DPCs were extracted and measured using the ARK assay. Asterisks indicate adjusted p-values: **p < 0.01, ***p < 0.001, ****p < 0.0001.

To investigate DPC formation, HeLa cells were treated with increasing concentrations of glyoxal and analyzed using the Advanced Recovery of K-SDS Precipitates (ARK) assay(31). Compared to the non-exposed cells, the 24-hour exposure at low micromolar concentrations resulted in DPC levels increasing by 1.20 ± 0.04-fold at 1 µM glyoxal to 1.46 ± 0.01-fold at 10 µM glyoxal (mean ± SD, n=4; **Figure 3B)**. Under 2-hour treatment conditions, DPC levels rose to 7.81 ± 0.74-fold, 16.84 ± 2.27-fold, and 24.64 ± 2.03-fold at 5, 10 and 20 mM glyoxal levels respectively (mean ± SD, n=3; **Figure 3C)**.

To confirm the ARK assay findings, DPCs were isolated from glyoxal treated HeLa cells using the Superior method for True DNA-protein crosslink Recovery (STAR) assay (32). Equal amounts of DNA were loaded and separated by SDS-PAGE. DNA-crosslinked proteins were visualized by silver staining. Glyoxal-treated samples exhibited a concentration-dependent increase in DNA-associated proteins compared with untreated control **(Figure 3D)**.

Because intracellular glyoxal concentrations are tightly regulated by the glyoxalase pathway, we next examined whether disruption of glyoxal detoxification pathway influences DPC formation. Wild type and GLO-2 knockout cells were treated with glyoxal, and DPC levels isolated and quantified with the ARK assay. While both cell lines exhibited increased DPC formation following glyoxal exposure, GLO-2 knockout cells accumulated a 1.57 ± 0.35-fold increase in DPCs when compared to wild-type cells **(Figure 3E)**. These findings indicate that efficient glyoxal detoxification limits glyoxal-induced formation of DNA-protein crosslinks. Collectively, these results demonstrate that glyoxal induces DNA-protein crosslinks in human cells and that the extent of DPC formation is influenced by intracellular glyoxal detoxification activity.

### SPRTN contributes to the repair of glyoxal-induced DNA-protein crosslinks

SPRTN is a specialized metalloprotease that plays crucial role in the proteolytic processing of DNA-protein crosslinks. To determine whether glyoxal-induced DPCs are repaired through this pathway, DPC accumulation was compared between control and SPRTN-knockdown cells following glyoxal treatment. Quantification using the ARK assay revealed SPRTN-deficient cells exhibited 1.51 ± 0.40-fold increase in DPC levels compared to wild-type controls following glyoxal exposure, indicating impaired removal of glyoxal-induced DPCs **(Figure 4A)**.

**Figure 4:**
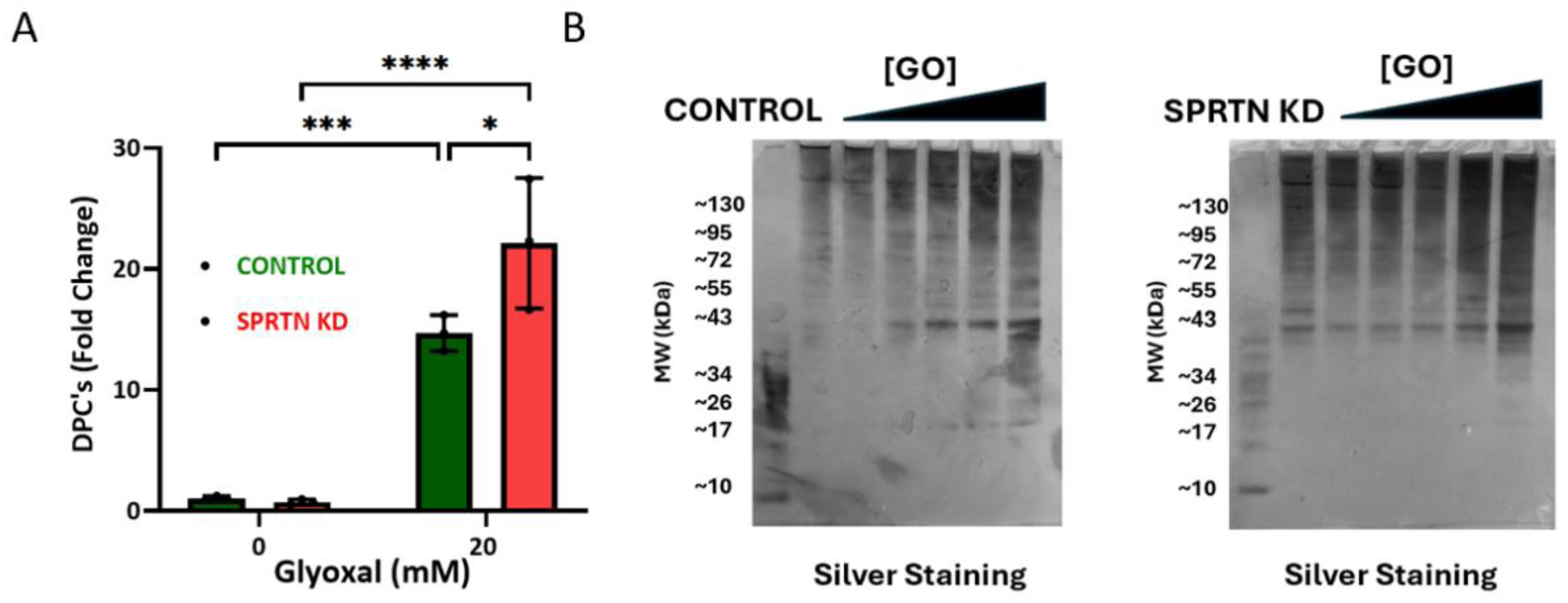
Effect of SPRTN on glyoxal-induced DPCs. A) DPC levels (shown as fold change) were measured in control and SPRTN-knockdown (SPRTN KD) cells treated with glyoxal (20 mM, 2 hours). DPCs were extracted and measured using the ARK assay. Asterisks indicate adjusted p-values: **p < 0.01, ***p < 0.001, ****p < 0.0001. B) SPRTN WT (SPRTN⁺^/^⁺) and SPRTN KD (SPRTN⁻^/^⁺) cells were treated with 0, 0.1, 1, 5, 10, or 20 mM GO for 2 hours and were processed using STAR assay to isolate DPCs. Isolated DPCs were resolved by 4-12%SDS-PAGE and visualized by the silver staining.

The elevated levels of DPCs in SPRTN-deficient cells were independently confirmed using the STAR assay. Consistent with the ARK results, glyoxal-treated SPRTN-deficient cells exhibited increased accumulation of DNA-crosslinked proteins compared with wild-type cells. These findings demonstrate that SPRTN contributes to the repair of glyoxal-induced DNA-protein crosslinks **(Figure 4B)**.

### Proteomic characterization of glyoxal-induced DNA-protein crosslinks

To identify proteins participating in glyoxal-induced DNA-protein crosslink formation we processed isolated DPCs by mass spectrometry-based proteomics. DPCs isolated using the STAR assay from control and glyoxal-treated HeLa cells (20 mM, 2 hours) were analyzed by silver staining of SDS-Page DPC protein separation and label-free LC-MS/MS (**Figure S2**). The reproducibility of biological replicates was assessed by Pearson correlation analysis using normalized log_2_-transformed protein abundance values (**Figure S3**). High correlations were observed within both the control (0.83 ± 0.04) and glyoxal-treated (0.79 ± 0.07) groups, whereas lower correlations were observed between control and glyoxal-treated samples (0.66 ± 0.06), indicating that glyoxal treatment produced reproducible changes in the DPC-associated proteome.

Differential protein enrichment analysis revealed substantial alterations in the DPC-associated proteome following glyoxal treatment. The results were visualized using a volcano plot of log_2_ fold change versus -log_10_(p-value), which revealed 469 proteins significantly enriched in glyoxal-treated samples compared with untreated controls. Among the enriched proteins were several chromatin-associated and DNA-binding proteins, including H2A, HMGN5, PCNA, and RPA3. These identifications suggest that glyoxal preferentially crosslinks proteins involved in chromatin organization, DNA replication, and DNA repair **(Figure 5A)**.

**Figure 5:**
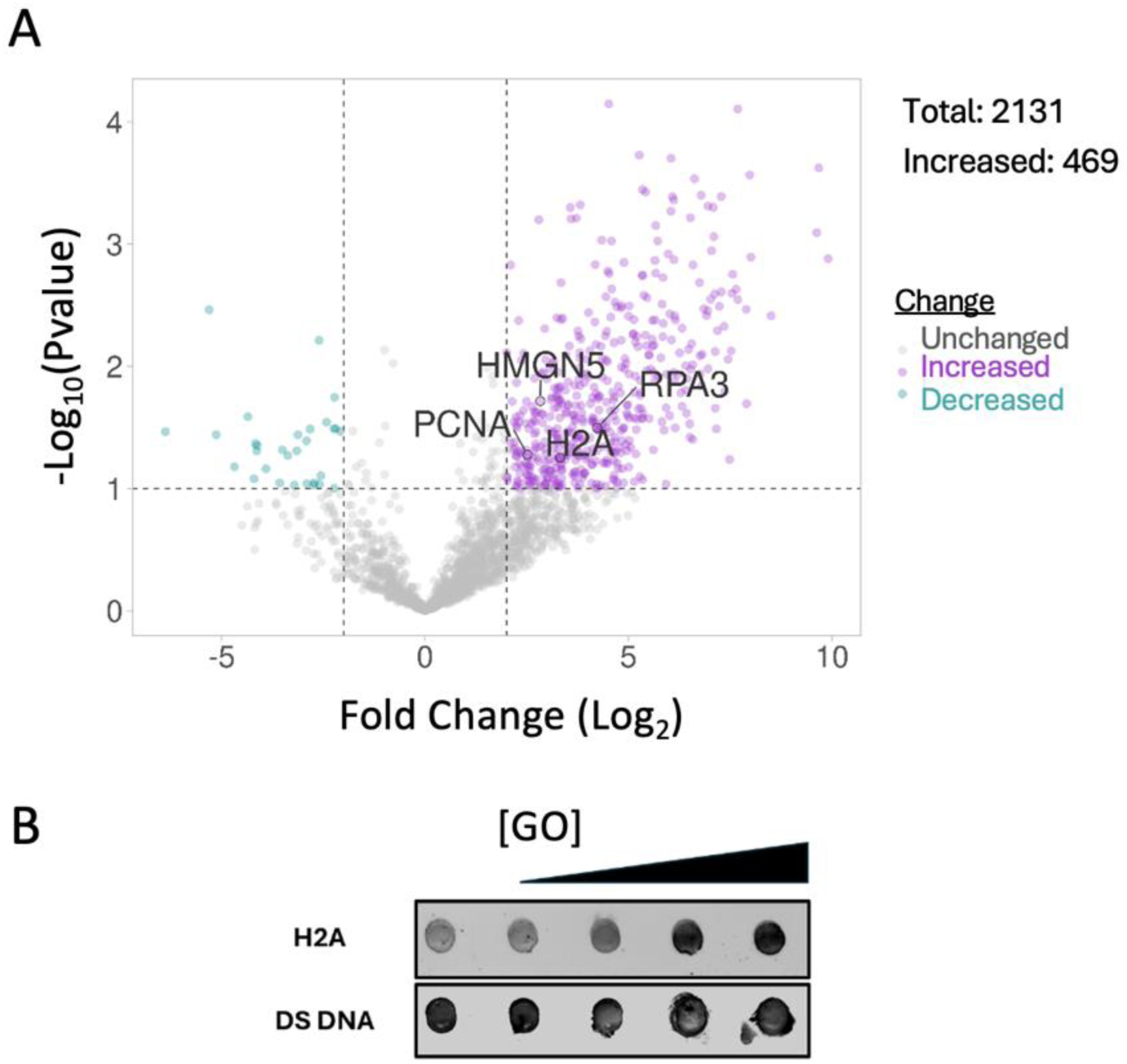
Proteomic results of glyoxal-induced DPCs. A) The plot shows fold change (log2) versus significance (-log10 pvalue). Proteins with increased abundance are shown in purple, decreased in blue, and unchanged in gray. Various enriched proteins (H2A, 49PCNA, RPA3, HMGN5) are highlighted. A total of 2131 proteins were identified, with 469 proteins showing increased levels. B) Representative dot blots of DPCs isolated via STAR assay from HeLa cells treated with 0, 0.1, 5,10, or 20 mM GO for 2 hours. Samples were normalized for DNA content, immobilized on nitrocellulose membranes, and probed with primary antibodies specific for histone H2A and dsDNA.

To confirm the change in the DPC-associated proteome following glyoxal treatment, we performed dot blot analysis to measure the change in histone DNA interaction. HeLa cells were treated with 0, 0.1, 5,10, or 20 mM glyoxal for 2 hours and DPCs were extracted with the STAR assay. Dot blot analysis showed that antibody signal for histone H2A increased in a concentration-dependent manner, indicating glyoxal-dependent DNA-protein crosslink (DPC) formation **(Figure 5B)**. The dsDNA was used as a loading control and remained relatively constant across samples.

Gene ontology enrichment analysis was performed to gain further insight into the biological functions of proteins enriched within glyoxal-induced DPCs. The Benjamini-adjusted pvalue was used to assess significance of enrichment. Cellular component analysis demonstrated significant enrichment of proteins localized to the nucleus (7×10^-^¹¹⁸), chromatin (2.5×10^-^³), nucleosomes (1.05×10^-^²), and other chromosome telomeric regions (4×10^-19^) **(Figure 6A)**. Molecular function analysis revealed enrichment of proteins involved in protein binding (8.29×10^-^⁶¹), nucleic acid binding (1.71 × 10^-^⁸), and RNA binding (1.83 × 10^-^¹⁵⁰) **(Figure 6B)**. Biological process analysis further showed significant enrichment of proteins associated with chromatin organization (2×10^-^ ¹⁵), DNA repair (4×10^-^⁶), DNA damage response (6×10^-^⁵), transcriptional regulation (8.7 ×10^-^³), RNA processing (5×10^-^⁶³), mRNA splicing (4×10^-^⁶⁵) (**Figure 6C)**.

**Figure 6:**
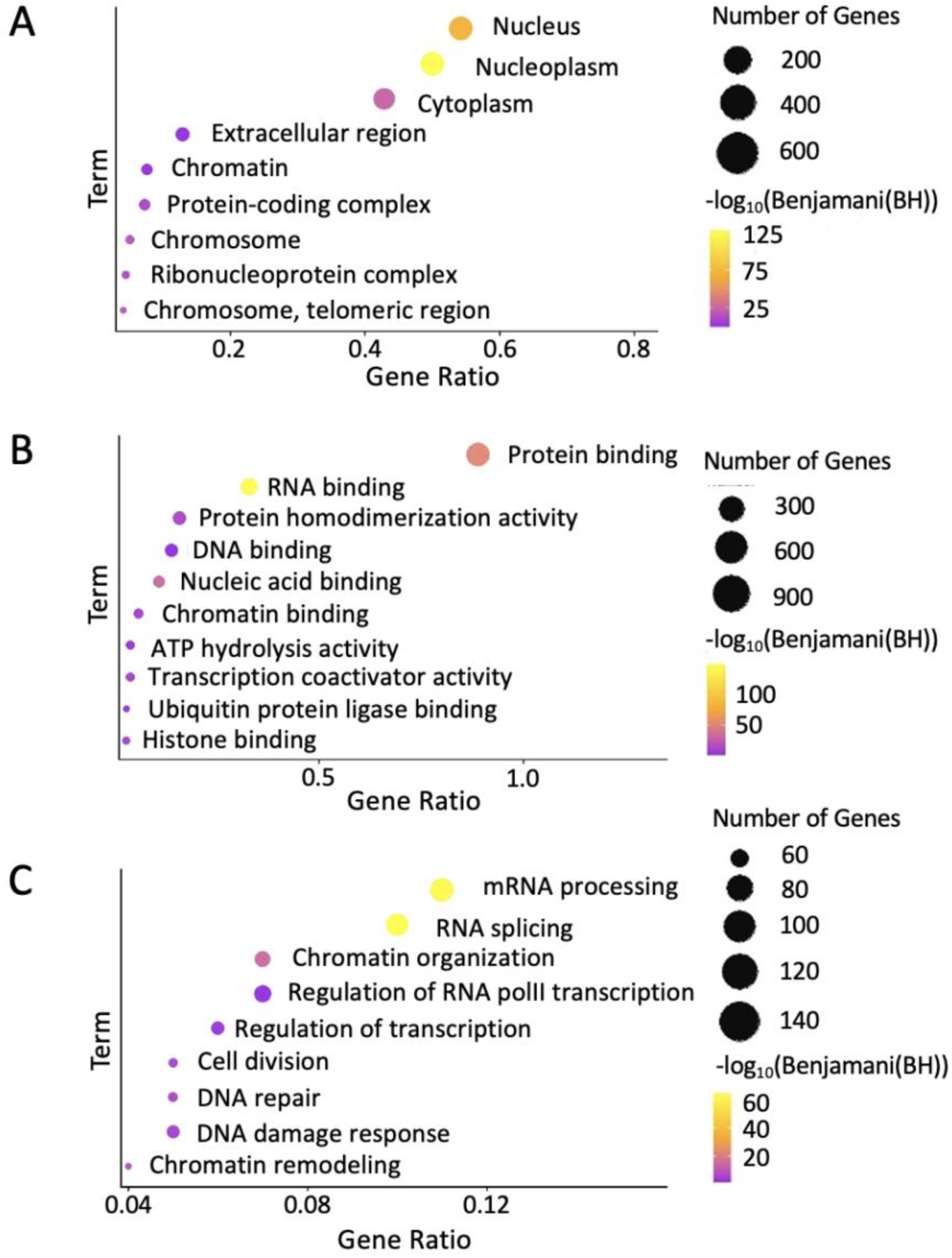
Gene ontology analysis of DPCs enriched by treatment of glyoxal. A) Cellular Compartment analysis was performed with the DAVID overrepresentation test, and results are shown as a multivariable plot with the -log10(BH), number of genes, and fold enrichment displayed for each enriched gene ontology cellular compartment term. B) Molecular function analysis was performed with the DAVID overrepresentation test, and results are shown as a multivariable plot with the -log10(BH), number of genes, and fold enrichment displayed for each enriched gene ontology molecular function term. C) Biological process analysis was performed with the DAVID overrepresentation test, and results are shown as a multivariable plot with the -log10(BH), number of genes, and fold enrichment displayed for each enriched gene ontology biological process term.

Collectively, these findings demonstrate that glyoxal induced DNA-protein crosslinks preferentially involve proteins responsible for maintaining chromatin organization, genome stability, and nucleic acid metabolism, providing proteome-wide insight into the cellular pathways susceptible to glyoxal-mediated DNA-protein crosslink formation.

To understand the biological relevance of glyoxal induced DPCs, we analyzed the subset of enriched proteins associated with DNA repair pathways and grouped them based on their functions. We found that several of the enriched proteins participate in DNA repair. Some proteins, such as RAD23A, RAD23B, CETN2, and RPA3, are part of the nucleotide excision repair (NER) pathway (36-39), which is a major repair pathway to remove bulky DNA modifications. We also identified PPP4R2, which is linked to the homologous recombination (HR) DNA repair pathway (40). In addition, proteins like NPM1, YWHAQ, SUMO2/3, NEDD8, PML are involved in the overall DNA damage response (DDR)(41-45), helping cells recognize and respond to DNA damage. ISY1, which is associated with base excision repair pathway (BER) (46), was also detected. These results show that many different DNA repair proteins exhibit elevated DNA-protein interactions, suggesting that glyoxal induced DNA damage is complex and requires multiple repair pathways.

### *In-vitro* confirmation of glyoxal-induced DNA-protein crosslink formation

Our proteomic analysis identified several DNA-binding and chromatin-associated proteins in glyoxal-induced DNA-protein crosslinks. Histone proteins are core proteins that package DNA into nucleosomes and have previously been reported to be susceptible to glyoxal modification(47). We therefore examined whether glyoxal could directly crosslink histone proteins to DNA. To determine whether glyoxal can directly form DNA-protein crosslinks, we performed *in-vitro* reactions using purified recombinant histone and 6-carboxyfluorescein-labeled DNA. Incubation of histone H2A and labeled-DNA sequence together with increasing concentrations of glyoxal (0, 0.5, 1, 5 and 10 mM) resulted in a new band at around 20 kDa. This band matches the combined molecular weight when H2A is crosslinked to the labeled DNA. Furthermore, this band was present at the same position of the SDS-page gel when visualized by silver staining and fluorescent imaging, indicating that a crosslink formed. The intensity of this band elevated as the glyoxal concentration increased, showing that crosslink formation is concentration dependent. Importantly, treatment with proteinase-K (PK) resulted in a loss of high molecular weight DNA signal in the fluorescent imaging **(Figure 7)**. Incubation of histones H2B, H3.1, and H4 with labeled DNA and glyoxal also formed crosslink products **(Figure S4-S6)**. In summary, these findings confirm histones are susceptible to formation of DNA-protein crosslinks upon glyoxal exposure.

**Figure 7:**
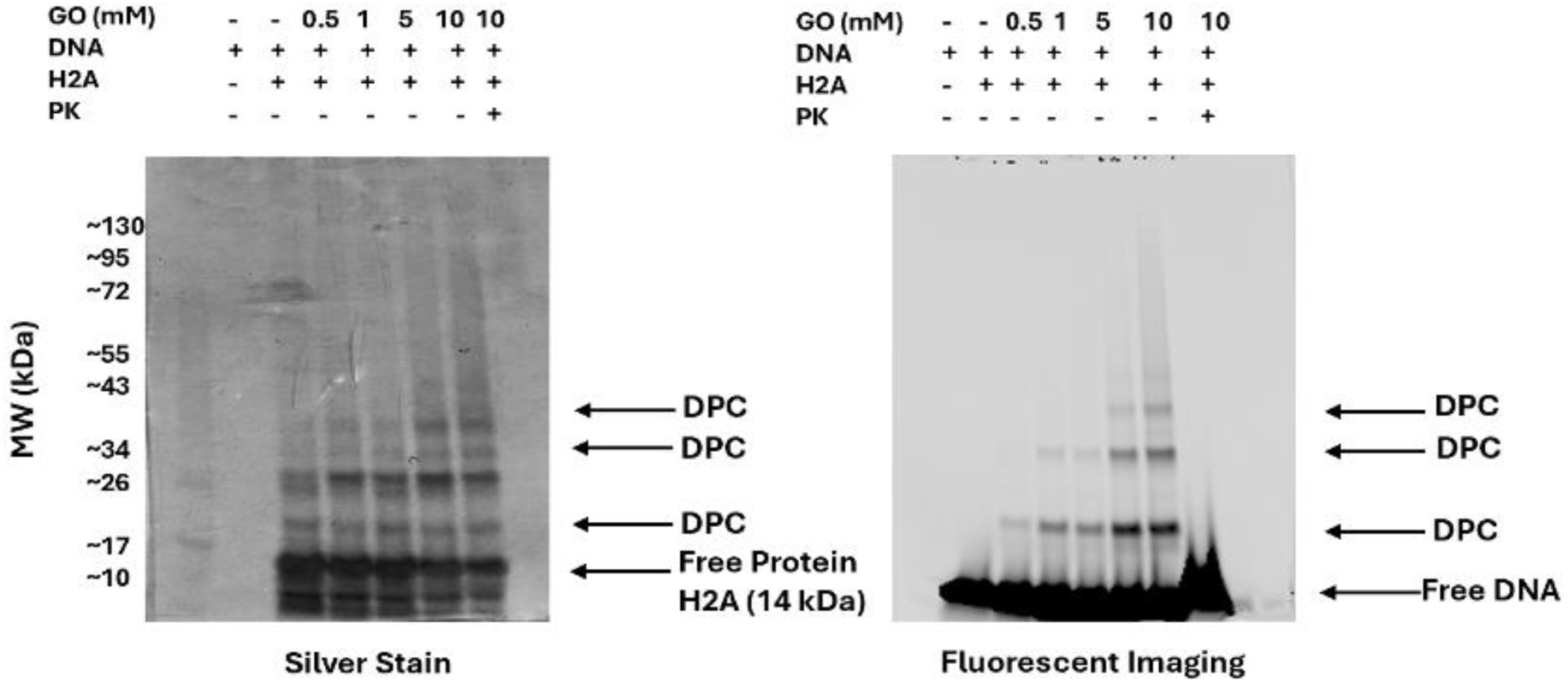
Glyoxal induces DNA-Protein crosslink (DPC) formation in vitro. A) Histone H2A (1 μg) was incubated with a fluorescently labelled DNA containing the sequence (TTAGGG)_3_ (1 μg) in the presence of increasing concentrations of glyoxal in PBS (pH 7.4) at 37°C for 2 h (total volume 25 μL). Samples were analyzed by SDS-PAGE followed by silver staining (left) and fluorescence imaging (right).

To understand the specificity of amino acids in crosslinking, we performed capping experiments. We blocked histone lysine residues using 2,5-dioxopyrrolidin-1-yl acetate (NHS-ester), cysteine residues using iodoacetamide (IA), and arginine using 1,2-cyclohexadione (CHD) before adding glyoxal. When histone lysine residues were blocked, the crosslink formation was clearly reduced, suggesting lysine plays an important role **(Figure S7)**. Blocking cysteine had minimal effect, indicating it is not a major contributor **(Figure S8)**. When both arginine and lysine were blocked, crosslink formation decreased further, suggesting that these amino acids, especially lysine and arginine participate in glyoxal induced crosslinking **(Figure S9)**.

## DISCUSSION

Defining which endogenous metabolites contribute to DNA-protein crosslinking is important for understanding their functional ramification and mutagenic potential (1). In this study, we use a variety of approaches to demonstrate that glyoxal induces DNA-protein crosslinks (DPCs) in human cells. Using ARK and STAR assays, we show that DPC levels increase significantly upon glyoxal exposure, consistent with previous reports that reactive carbonyl species can induce covalent DNA-protein linkages(13, 14, 48, 49). Baseline levels of DPCs observed in untreated cells likely arise from endogenous metabolites such as reactive oxygen species (50), methylglyoxal (51) and formaldehyde(2, 49), as well as normal cellular processes involving transient protein-DNA interactions(1, 2).

Mechanistically, glyoxal is a highly reactive dicarbonyl that modifies nucleophilic residues such as lysine and arginine in proteins and guanine bases in DNA. Our *in-vitro* experiments with histone proteins support a mechanism in which glyoxal covalently links DNA and proteins. This is consistent with previous studies that reactive aldehydes such as formaldehyde, which can form reversible Schiff base intermediates that stabilize into covalent DNA-Protein crosslinks (52). From a biological perspective, glyoxal is produced through glucose autooxidation, lipid peroxidation, and metabolic pathways(15, 16, 53). Glyoxal levels are elevated in conditions such as diabetes and neurodegeneration (24, 54, 55). The ability of glyoxal to induce DPCs provides a potential mechanistic link between metabolic imbalance and DNA damage. Therefore, glyoxal-induced DPC formation may contribute to genomic stress (56) and cellular dysfunction under conditions characterized by elevated reactive carbonyl species.

Proteomic analysis revealed that glyoxal-induced DPCs involve a broad range of nuclear proteins, particularly those associated with chromatin organization, transcription, DNA replication and repair. Multiple chromatin-associated proteins were increased, including histone proteins (H2A, H4C1), chromobox proteins (CBX1, CBX3, and CBX5), chromatin remodeling factors such as SMARCE1 and RBBP4. In addition, several transcription-associated proteins and RNA polymerase subunits, including POLR2F, POLR2I, POLR2E, and POLR2H, were identified, suggesting that glyoxal-induced crosslinking may interfere with transcriptional machinery. Proteins involved in DNA replication and genome maintenance, including PCNA, ORC2, GINS4, POLE3, RPA3, TP53BP1, RAD23A, and RAD23B were also increased, supporting the idea that glyoxal-induced DPCs may disrupt DNA replication and DNA damage response pathways. Furthermore, the enrichment of RNA-binding and RNA-processing proteins such as HNRNPA1, HNRNPA2B1, SRSF family proteins, and LSM complex proteins suggests that glyoxal exposure may broadly affect nucleic acid-associated protein networks. Similar proteomic results have been reported for methylglyoxal-induced DPCs, where hundreds of proteins involved in nucleosome assembly, transcription, and RNA processing were identified (14). Collectively, these findings suggest that glyoxal -induced crosslinking may disrupt essential processes such as DNA replication and transcription, thereby contributing to genomic instability.

Our results also highlight the vital role of SPRTN in repairing glyoxal-induced DPCs. We observed increased accumulation of DPCs in SPRTN-deficient cells, indicating that SPRTN contributes to efficient removal of these lesions. SPRTN is a DNA-dependent metalloprotease that degrades proteins crosslinked to DNA during replication(6, 57). This finding is consistent with previous reports showing that loss of SPRTN leads to accumulation of DPCs, replication stress and genomic instability as observed in Ruijs Aalfs syndrome (3). In addition to SPRTN, proteasome-mediated degradation (58, 59) and p97-dependent pathways(60-62) have also been shown to contribute to DPC repair, highlighting the key involvement of proteolytic systems.

Post-translational modifications such as ubiquitination and SUMOylation also appear to play important roles in regulating DPC repair. The enrichment of SUMO 2/3 and NEDD8 in our dataset suggests that protein modification pathways may facilitate recognition and processing of crosslinked proteins. These modifications are known to regulate protein stability, localization, and interactions, and have been involved in coordinating DNA damage responses and repair processes(43, 44, 63). However, the enrichment of SUMO 2/3 and NEDD8 observed does not directly establish a functional role in glyoxal-induced DPC repair, and further mechanistic studies will be necessary to determine whether these pathways actively participate in DPC recognition and processing.

Despite these insights, this study has limitations. Proteomic analysis identifies the proteins participating in DPC formation, but these observations do not provide information about exact chemical structures or crosslinking sites. In addition, although ARK and STAR assays enrich DNA-associated protein complexes, these methods cannot completely distinguish direct covalent DNA-protein crosslinks from tightly bound non-covalent protein interactions that may persist during isolation (64). Future studies can include structural characterization of glyoxal-induced DPCs, identification of specific amino acid-DNA linkages, and site-specific mapping of crosslinks using high-resolution mass spectrometry approaches. Experiments to develop analytical methods to uncover the amino acids participating in glyoxal DNA cross-linking and conduct glyoxal DPC characterization experiments under physiological levels of glyoxal concentrations are underway.

In conclusion, our study demonstrates that glyoxal is a potent inducer of DPCs in living cells, targeting a wide range of chromatin-associated proteins and engaging multiple DNA repair pathways. These findings expand our understanding of how endogenous reactive metabolites contribute to DNA damage and highlight the potential role of glyoxal-induced DPCs in disease.

## ACKNOWLEDGEMENTS

We thank members of the Erber lab, Alexander Hurben, Steven Bloom and Mark Farrell for their insightful suggestions and discussion. Support for this work was provided by startup funds from the University of Kansas (PI: Erber, L) and funded by a grant from NIGMS of the National Institutes of Health (R35GM159935, PI: Erber, L).

## NOTES

The authors declare no competing financial conflict.

