## Supplemental Information for "Glyoxal induces DNA-Protein Crosslinking in Cells"

**Table of Contents**


I. Supplementary Figures Page 3
II. Supplementary Table Page 7

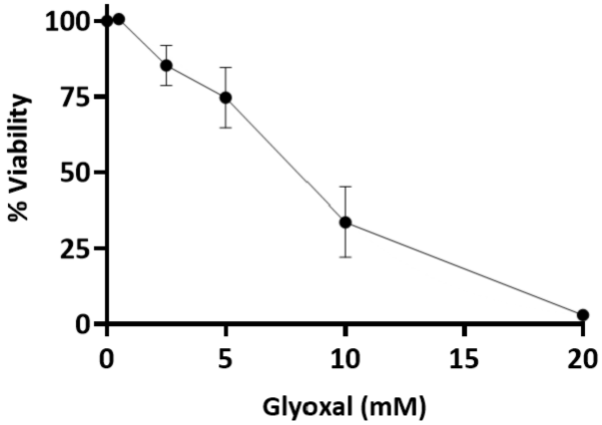


**Figure S1.** HeLa cells were treated with different concentrations of glyoxal (0, 0.5, 2.5, 5, 10 and 20 mM) for 6 h and cell viability was measured using the PrestoBlue assay. Data represents the mean ± SD of three independent replicates (n=3). Nonlinear regression analysis was performed using GraphPad Prism.

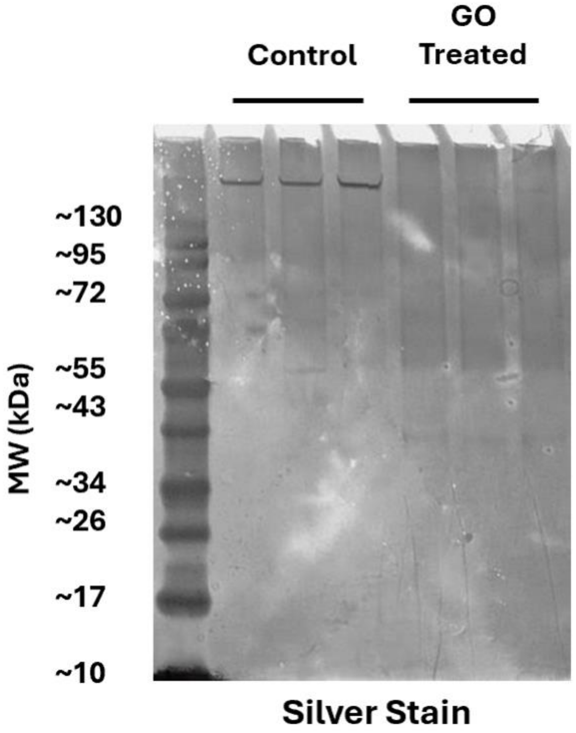


**Figure S2.** HeLa cells were treated with 20 mM glyoxal (GO) for 2 h. DNA–protein crosslinks (DPCs) were isolated using the STAR assay, separated on 4–12% SDS-PAGE gels, and visualized by silver staining.

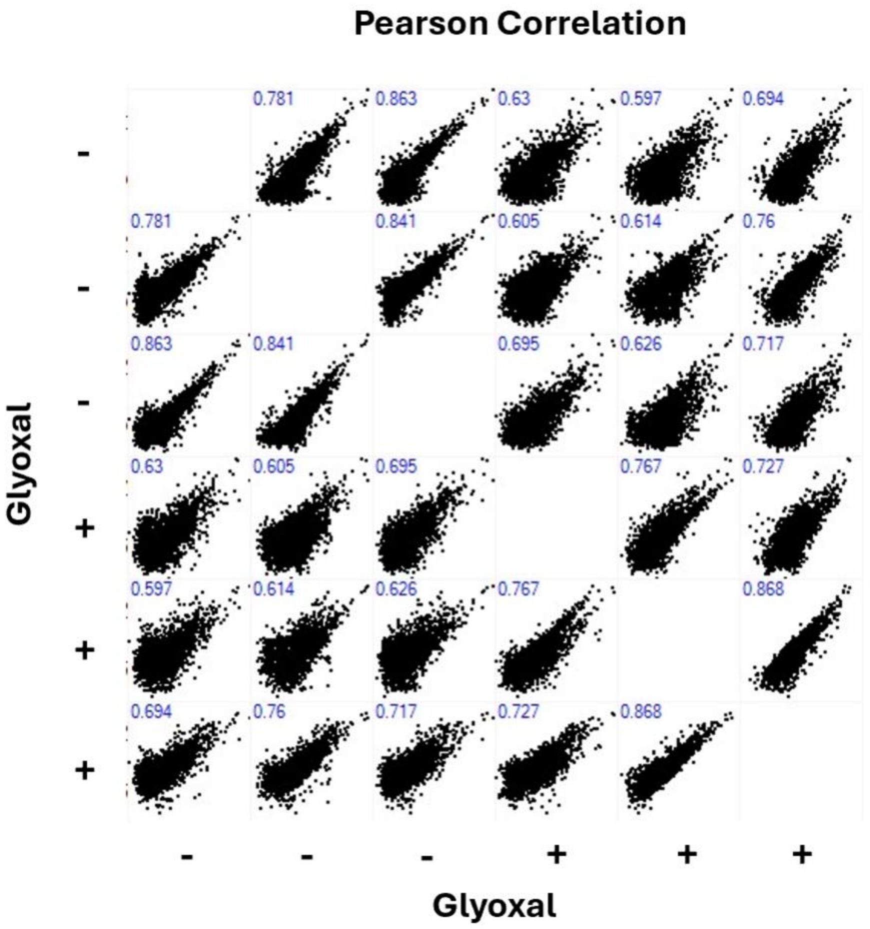


**Figure S3.** Pearson correlation analysis of normalized $log_2$-transformed protein abundance values obtained by label-free LC-MS/MS to assess reproducibility among biological replicates. Negative mark indicates control treatment. Positive mark indicates glyoxal treated (20 mM, 2 h).

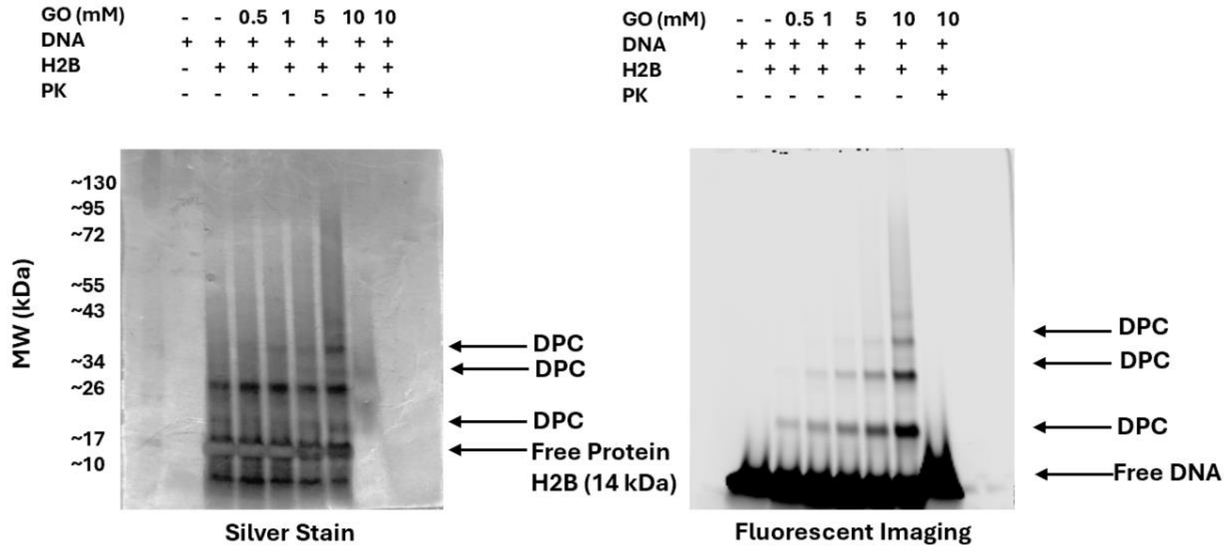


**Figure S4**: Glyoxal induces DNA-Protein crosslink (DPC) formation in vitro.
Histone H2B (1 µg) was incubated with a fluorescently labelled DNA containing the sequence $(TTAGGG)_3$ (1 µg) in the presence of increasing concentrations of glyoxal in PBS (pH 7.4) at 37$^{o}$C for 2 h (total volume 25 µL). Samples were analyzed by SDS-PAGE followed by silver staining (left) and fluorescence imaging (right).

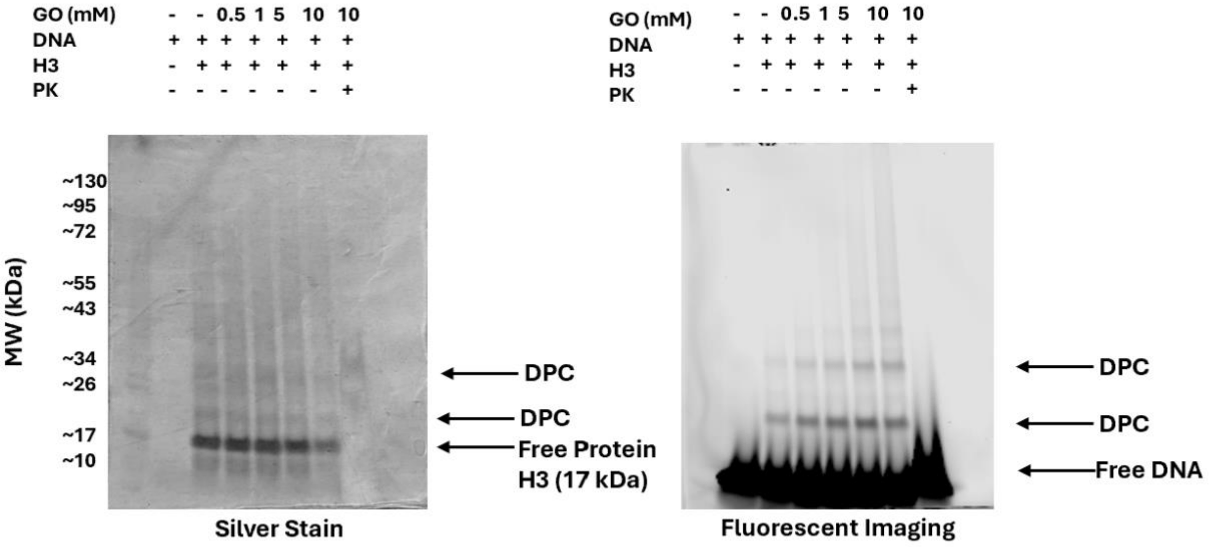


**Figure S5**: Glyoxal induces DNA-Protein crosslink (DPC) formation in vitro.
Histone H3 (1 µg) was incubated with a fluorescently labelled DNA containing the sequence $(TTAGGG)_3$ (1 µg) in the presence of increasing concentrations of glyoxal in PBS (pH 7.4) at 37$^{o}$C for 2 h (total volume 25 µL). Samples were analyzed by SDS-PAGE followed by silver staining (left) and fluorescence imaging (right).

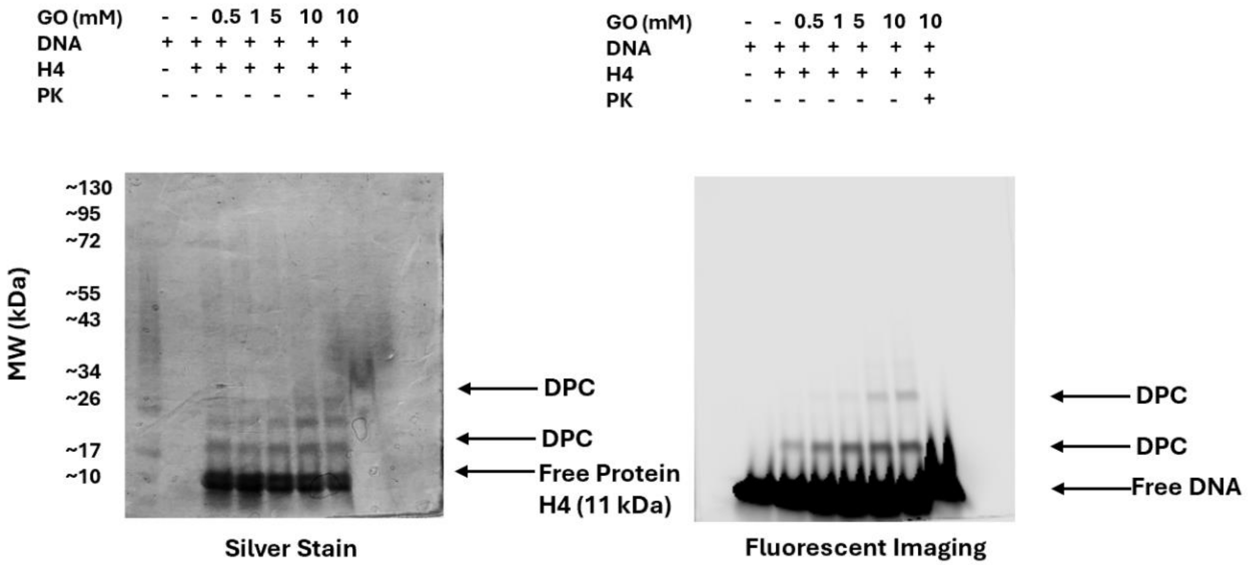


**Figure S6**: Glyoxal induces DNA-Protein crosslink (DPC) formation in vitro.

Histone H4 (1 µg) was incubated with a fluorescently labelled DNA containing the sequence $(TTAGGG)_3$ (1 µg) in the presence of increasing concentrations of glyoxal in PBS (pH 7.4) at 37°C for 2 h (total volume 25 µL). Samples were analyzed by SDS-PAGE followed by silver staining (left) and fluorescence imaging (right).

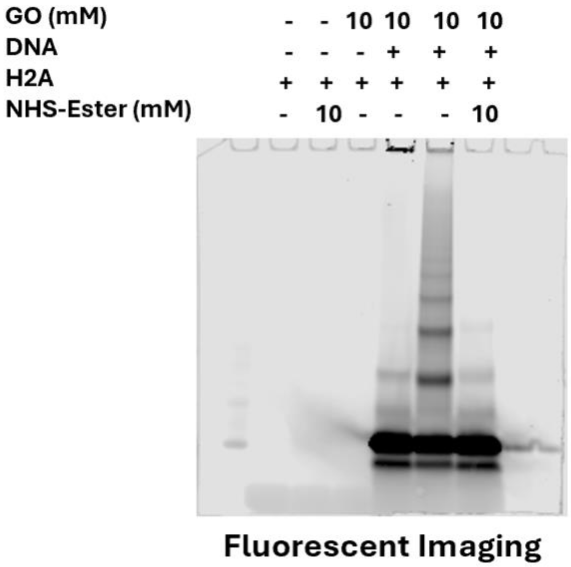


**Figure S7.** Amino-acid capping experiment was performed by treating Histone H2A and 6-carboxyfluorescein-labeled DNA with glyoxal following chemical modification of lysine with 2,5-dioxopyrrolidin-1-yl acetate (NHS-ester) for 1h at 37°C. Samples were separated by SDS-Page and visualized by fluorescence imaging.

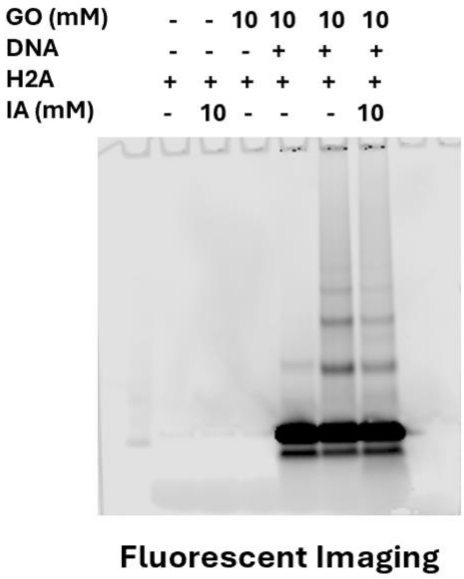


**Figure S8.** Amino-acid capping experiment was performed by treating Histone H2A and 6-carboxyfluorescein-labeled DNA with glyoxal following chemical modification of cysteine with iodoacetamide for 1h at room temperature in the dark. Samples were separated by SDS-Page and visualized by fluorescence imaging.

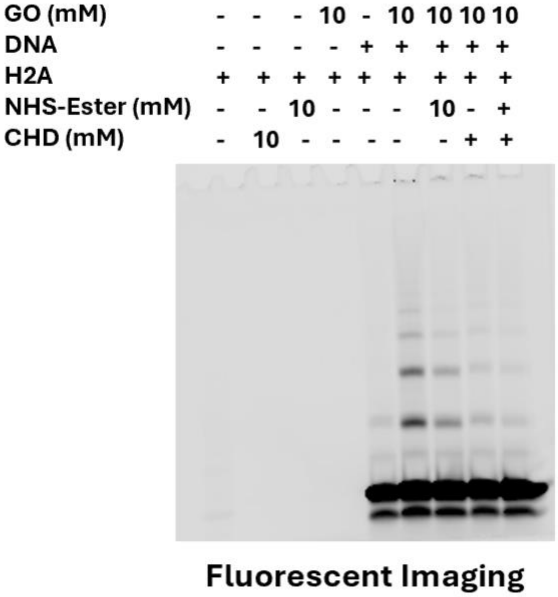


**Figure S9.** Amino-acid capping experiment was performed by treating Histone H2A and 6-carboxyfluorescein-labeled DNA with glyoxal following chemical modification of lysine with 2,5-dioxopyrrolidin-1-yl acetate (NHS-ester) and arginine with 1,2-cyclohexadione (CHD) for 1h at 37°C. Samples were separated by SDS-Page and visualized by fluorescence imaging.

**Supplemental Table 1:** The table lists proteins identified in glyoxal-induced DPC samples, along with their UniProt Accession IDs, fold change ($log_2$) between glyoxal and vehicle control treatment groups, and statistical significance (-$log_{10}$(p-value)).

| Gene Name | Uniprot Accession ID | Fold change (log2) | Significance (-$log_{10}$(p-value)) |
|---|---|---|---|
| POLR2F | P61218 | 9.6720 | 3.6244 |
| LSM3 | P62310 | 9.8982 | 2.8809 |
| CDKN2A | P42771 | 9.6217 | 3.0938 |
| DPY30 | Q9C005 | 7.6791 | 4.1055 |
| SH3BGRL | Q9H299 | 7.9759 | 3.5663 |
| SET | Q01105 | 8.5001 | 2.4131 |
| LSM8 | O95777 | 8.0036 | 2.8944 |
| SMAP | O00193 | 7.2747 | 3.3893 |
| CHMP4B | Q9H444 | 7.0766 | 3.2996 |
| TPM1 | A0A494BZZ2 | 7.8883 | 2.4664 |
| ATOX1 | E5RIM7 | 7.5571 | 2.7524 |
| PSMD9 | O00233 | 7.6395 | 2.6300 |
| ERH | P84090 | 6.9463 | 3.3113 |
| PHPT1 | Q9NRX4 | 7.6808 | 2.5465 |
| UBE2R2 | Q712K3 | 6.7805 | 3.4008 |
| HSBP1 | O75506 | 7.0928 | 3.0613 |
| B4DLN1_HUMAN | B4DLN1 | 6.6169 | 3.5347 |
| RRP15 | Q9Y3B9 | 7.5380 | 2.5968 |
| S100A13 | Q99584 | 7.0371 | 2.9466 |
| CLTA | P09496 | 7.2759 | 2.6095 |
| NDUFAB1 | O14561 | 6.0413 | 3.7027 |
| JPT1 | Q9UK76 | 6.5172 | 3.2156 |
| CNDP2 | Q96KP4 | 7.2275 | 2.4912 |
| MYL9 | P24844 | 7.1150 | 2.5921 |
| CHRAC1 | Q9NRG0 | 7.5211 | 2.1267 |
| POLR2I | P36954 | 7.0130 | 2.5480 |
| CBX1 | P83916 | 7.4442 | 2.0566 |
| UTP20 | O75691 | 7.1818 | 2.2915 |
| CHCHD4 | Q8N4Q1 | 6.1084 | 3.3606 |
| PPP1R14B | Q96C90 | 6.0753 | 3.3892 |
| UFM1 | P61960 | 6.7435 | 2.6860 |
| HEXIM1 | O94992 | 6.5828 | 2.8291 |
| NPM3 | O75607 | 6.9430 | 2.4543 |
| EIF4EBP1 | Q13541 | 6.9108 | 2.4152 |
| CHMP4A | Q9BY43 | 6.8471 | 2.4703 |
| ARK2N | Q96B23 | 6.0406 | 3.2699 |
| PFDN6 | O15212 | 7.1249 | 2.1211 |
| GBA1 | P04062 | 7.3553 | 1.7929 |

| | | | |
|---|---|---|---|
| BPIFB4 | P59827 | 5.2590 | 3.7283 |
| ANAPC13 | Q9BS18 | 6.1128 | 2.8724 |
| PTMS | A0AA34QVV1 | 6.9532 | 1.9810 |
| TPM2 | Q5TCU3 | 6.8051 | 2.0972 |
| POLR1D | P0DPB6 | 5.9705 | 2.9196 |
| S100A10 | P60903 | 5.4147 | 3.4275 |
| ASDURF | L0R819 | 6.7554 | 2.0862 |
| ZNF428 | M0QXZ5 | 5.6673 | 3.1515 |
| BASP1 | P80723 | 7.1288 | 1.6573 |
| SMPD3 | Q9NY59 | 5.3393 | 3.4466 |
| RBM8A | Q9Y5S9 | 5.7102 | 3.0250 |
| PAWR | Q96IZ0 | 6.5466 | 2.1849 |
| MIF | P14174 | 7.4816 | 1.2374 |
| NES | P48681 | 6.7604 | 1.9387 |
| CHMP2A | M0R1T5 | 4.5107 | 4.1476 |
| C1QBP | Q07021 | 7.1555 | 1.4874 |
| GRB2 | P62993 | 6.7932 | 1.8400 |
| TBCA | E5RJD8 | 6.3135 | 2.3124 |
| TPM4 | P67936 | 6.4733 | 2.1382 |
| FIP1L1 | A0A994J6B4 | 6.5778 | 2.0135 |
| PFDN4 | Q9NQP4 | 5.8478 | 2.7262 |
| HYPK | Q9NX55 | 6.1493 | 2.3789 |
| CHTF18 | A0A0D9SF58 | 5.6469 | 2.8664 |
| TADA3 | O75528 | 6.5205 | 1.9750 |
| NOL3 | O60936 | 6.0138 | 2.4280 |
| HDGF | A0AA34QVG5 | 5.6684 | 2.7614 |
| DUT | P33316 | 6.4053 | 2.0032 |
| LEO1 | Q8WVC0 | 6.2518 | 2.1112 |
| COX17 | Q14061 | 5.8245 | 2.5080 |
| GLO1 | Q04760 | 6.4339 | 1.8986 |
| PPP4R2 | A0AA34QVI2 | 6.0526 | 2.2163 |
| NOP16 | A0A0C4DGU5 | 6.4114 | 1.8058 |
| CRIP2 | P52943 | 6.2034 | 1.9327 |
| COPRS | Q9NQ92 | 5.3541 | 2.7452 |
| CBX3 | Q13185 | 5.8371 | 2.2428 |
| SH3BGRL3 | Q9H299 | 5.3291 | 2.7419 |
| WDR5 | P61964 | 6.2850 | 1.7833 |
| Krt18 | P05784 | 6.1061 | 1.9501 |
| PDCD5 | O14737 | 6.4252 | 1.6156 |
| CLTB | P09497 | 6.5013 | 1.5291 |
| MYL12A | J3QRS3 | 5.4533 | 2.5618 |
| ATP5F1D | P30049 | 6.1360 | 1.8775 |
| ANP32E | Q9BTT0 | 6.3075 | 1.6724 |

| | | | |
|---|---|---|---|
| PDAP1 | Q13442 | 6.5753 | 1.4016 |
| MYL6 | G8JLA2 | 5.5883 | 2.3753 |
| NPM1 | A0A7I2V3U2 | 5.3752 | 2.5818 |
| DFFA | O00273 | 6.5059 | 1.4177 |
| POLE3 | Q9NRF9 | 5.9321 | 1.9352 |
| OGFR | Q9NZT2 | 5.7888 | 2.0695 |
| CSNK2B | A0A0G2JM12 | 5.5822 | 2.2198 |
| RAD23B | P54727 | 5.6245 | 2.1723 |
| PCNP | Q8WW12 | 5.5853 | 2.1648 |
| WDR44 | Q5JSH3 | 4.8845 | 2.8491 |
| ANP32A | P39687 | 5.4138 | 2.2550 |
| SRSF9 | Q13242 | 5.7369 | 1.9119 |
| CDC26 | Q8NHZ8 | 6.1452 | 1.4962 |
| WBP11 | Q9Y2W2 | 6.0106 | 1.5950 |
| MLLT11 | Q13015 | 4.5790 | 3.0257 |
| EDF1 | O60869 | 5.4341 | 2.1691 |
| FLYWCH2 | I3L1Y9 | 5.5029 | 2.0962 |
| SMARCE1 | Q969G3 | 5.4971 | 2.0906 |
| MRFAP1 | Q9Y605 | 5.6142 | 1.9489 |
| LAMB1 | P07942 | 5.6493 | 1.8814 |
| GCSH | P23434 | 5.5452 | 1.9759 |
| CSTB | P04080 | 5.0166 | 2.4947 |
| CRIM1 | Q9NZV1 | 5.1282 | 2.3823 |
| CKS2 | P33552 | 5.2065 | 2.2436 |
| RGCC | Q9H4X1 | 5.4468 | 1.9636 |
| HTATSF1 | O43719 | 4.9977 | 2.4113 |
| CETN2 | P41208 | 5.2274 | 2.1607 |
| MT2A | P02795 | 5.1143 | 2.2628 |
| CALM2 | E7EMB3 | 4.3383 | 3.0336 |
| DDT | J3KQ18 | 5.1088 | 2.2621 |
| TPM3 | A0A087WWU8 | 5.1659 | 2.1978 |
| NHP2 | Q9NX24 | 5.8147 | 1.5472 |
| SNRPG | C9JVQ0 | 5.8698 | 1.4779 |
| TPT1 | A0A0B4J2C3 | 4.8233 | 2.4964 |
| HSPE1 | P61604 | 4.8174 | 2.4781 |
| UQCRH | P07919 | 4.5202 | 2.7594 |
| CREG1 | A0A3B3IRL2 | 4.4905 | 2.7793 |
| PTMA | H7C2N1 | 4.5815 | 2.6340 |
| PFDN1 | O60925 | 4.7100 | 2.4871 |
| DNAJC8 | O75937 | 4.6633 | 2.5260 |
| SRP9 | P49458 | 5.2879 | 1.8842 |
| TMA7B | A0A024R1R8 | 3.8092 | 3.3218 |
| RAD23A | P54725 | 5.1671 | 1.9201 |

| | | | |
|---|---|---|---|
| PEA15 | Q15121 | 5.5079 | 1.5728 |
| MPHOSPH6 | Q99547 | 5.2922 | 1.7868 |
| ARPP19 | H3BMD8 | 5.2849 | 1.7627 |
| DR1 | Q01658 | 4.8186 | 2.2138 |
| MYL6B | A0A8Q3SIC5 | 4.4413 | 2.5686 |
| MRPL49 | Q13405 | 4.1779 | 2.8319 |
| CCDC50 | Q8IVM0 | 4.9120 | 2.0622 |
| LONP1 | P36776 | 5.3523 | 1.6139 |
| AKAP8 | O43823 | 5.3088 | 1.6555 |
| RWDD1 | Q9H446 | 4.5612 | 2.4029 |
| BTF3L4 | Q96K17 | 4.9374 | 2.0244 |
| FUBP1 | A0A994J3Q8 | 5.9201 | 1.0370 |
| CHMP2B | A0A087WW88 | 5.0084 | 1.9352 |
| EEF1D | E9PL71 | 5.3516 | 1.5787 |
| FH | P07954 | 3.7135 | 3.2126 |
| SAFB | Q15424 | 5.6071 | 1.3165 |
| SNRPB2 | P08579 | 5.1274 | 1.7857 |
| SP100 | P23497 | 5.3264 | 1.5717 |
| CDH2 | P19022 | 5.2356 | 1.6520 |
| BCL7C | I3L1Q2 | 4.8995 | 1.9811 |
| LSM6 | P62312 | 3.5673 | 3.2987 |
| RANBP1 | F6WQW2 | 4.6402 | 2.1930 |
| CHMP3 | Q9Y3E7 | 4.8203 | 1.9943 |
| TXN | P10599 | 3.5870 | 3.2067 |
| CHMP1A | F5H875 | 4.4640 | 2.2865 |
| HEBP2 | Q9Y5Z4 | 5.1409 | 1.5799 |
| SAMD13 | H7BZX5 | 4.4269 | 2.2479 |
| CHMP1B | Q7LBR1 | 5.1873 | 1.4821 |
| PAIP2 | Q9BPZ3 | 4.9818 | 1.6839 |
| BAG2 | O95816 | 4.9571 | 1.6827 |
| ABRACL | Q9P1F3 | 4.7530 | 1.8364 |
| SELENOH | Q8IZQ5 | 5.0122 | 1.5756 |
| GSR | P00390 | 4.8027 | 1.7827 |
| CALU | O43852 | 4.9591 | 1.6147 |
| NACA | E9PAV3 | 4.9738 | 1.5948 |
| EMD | P50402 | 4.1701 | 2.3635 |
| RPLP2 | P05387 | 4.4711 | 2.0363 |
| SAP18 | O00422 | 4.4447 | 2.0525 |
| EXOSC6 | Q5RKV6 | 4.3196 | 2.1646 |
| VBP1 | P61758 | 5.1794 | 1.2841 |
| LSM7 | Q9UK45 | 4.0142 | 2.4394 |
| TERF2IP | Q9NYB0 | 5.3415 | 1.0719 |
| NIT2 | Q9NQR4 | 4.3470 | 2.0609 |

| | | | |
|---|---|---|---|
| KRT15 | P19012 | 5.0133 | 1.3930 |
| RBM17 | Q96I25 | 4.6088 | 1.7890 |
| POLR2H | P52434 | 4.6427 | 1.7488 |
| DAZAP1 | Q96EP5 | 5.0460 | 1.3382 |
| CIAO2B | Q9Y3D0 | 4.8825 | 1.4977 |
| SLTM | Q9NWH9 | 5.2783 | 1.0960 |
| TAF10 | Q12962 | 4.1949 | 2.1788 |
| BCL7B | Q9BQE9 | 4.4772 | 1.8813 |
| SNCG | O76070 | 4.8646 | 1.4924 |
| LRRFIP2 | Q9Y608 | 4.6282 | 1.7225 |
| GON7 | Q9BXV9 | 4.6595 | 1.6449 |
| NUP62 | P37198 | 4.9370 | 1.3595 |
| SNRPA1 | P09661 | 4.8189 | 1.4746 |
| RCN2 | Q14257 | 4.9646 | 1.3095 |
| SSB | P05455 | 4.3480 | 1.9132 |
| CCS | O14618 | 4.6943 | 1.5534 |
| PHAX | Q9H814 | 4.8380 | 1.3920 |
| LAMTOR5 | A0A8Z5A536 | 4.6263 | 1.5939 |
| CTNNA1 | P35221 | 5.1380 | 1.0555 |
| PFDN2 | Q9UHV9 | 4.5513 | 1.6366 |
| POLDIP3 | F6VRR5 | 5.0013 | 1.1757 |
| HNRNPDL | O14979 | 4.0466 | 2.1292 |
| CFDP1 | Q9UEE9 | 4.4543 | 1.7079 |
| SAP30BP | J3QQJ0 | 4.7819 | 1.3349 |
| PELP1 | C9JFV4 | 4.8119 | 1.2943 |
| KRT13 | P13646 | 4.6707 | 1.4188 |
| PPIL1 | Q9Y3C6 | 3.8276 | 2.2250 |
| PPIA | P62937 | 3.7570 | 2.2889 |
| NUP50 | Q9UKX7 | 4.6581 | 1.3834 |
| EEF1B2 | P24534 | 4.2805 | 1.7591 |
| DBI | B8ZWD1 | 3.3389 | 2.6849 |
| C19orf53 | Q9UNZ5 | 4.2739 | 1.7498 |
| ABI2 | F8WAL6 | 3.7710 | 2.2487 |
| ANP32B | Q92688 | 3.9590 | 2.0567 |
| ACAP1 | Q15027 | 4.4651 | 1.5455 |
| TMOD3 | Q9NYL9 | 4.5425 | 1.4607 |
| WDR12 | Q9GZL7 | 4.8373 | 1.1618 |
| SOD1 | P00441 | 2.7935 | 3.2001 |
| CAPZA1 | P52907 | 4.6811 | 1.3028 |
| PAGE1 | O75459 | 4.2027 | 1.7774 |
| YWHAB | P31946 | 4.1499 | 1.8298 |
| PPP2CA | A0A8V8TRB3 | 4.2332 | 1.7332 |
| CDV3 | Q9UKY7 | 4.4517 | 1.4977 |

| | | | |
|---|---|---|---|
| PGM2 | Q96G03 | 4.6504 | 1.2873 |
| NEDD8 | Q15843 | 4.5643 | 1.3725 |
| ARPC5 | O15511 | 4.0168 | 1.9138 |
| POLR2E | P19388 | 4.7180 | 1.2091 |
| NHERF1 | O14745 | 4.8799 | 1.0462 |
| GPI | A0A2U3TZU2 | 4.7474 | 1.1677 |
| ACOX1 | Q15067 | 3.9752 | 1.9385 |
| CACNA1E | A0A8V8TPU6 | 4.2708 | 1.6422 |
| PIN4 | Q9Y237 | 3.7987 | 2.1113 |
| LZIC | Q8WZA0 | 4.5995 | 1.3050 |
| MTPN | P58546 | 3.8054 | 2.0665 |
| SPRR2G | Q9BYE4 | 4.5596 | 1.3055 |
| CETN3 | E5RJF8 | 4.6330 | 1.2277 |
| LASP1 | Q14847 | 4.3771 | 1.4807 |
| ALYREF | E9PB61 | 4.1113 | 1.7457 |
| LAMP2 | A0A9L9PXQ4 | 3.8073 | 1.9892 |
| PSME3IP1 | Q9GZU8 | 4.2997 | 1.4903 |
| EIF1AX | P47813 | 3.8513 | 1.9384 |
| ELOB | Q15370 | 3.9140 | 1.8672 |
| GSTP1 | P09211 | 3.3884 | 2.3921 |
| CKS1B | P61024 | 4.7164 | 1.0524 |
| ISY1 | Q9ULR0 | 4.0610 | 1.6941 |
| ULK4 | Q96C45 | 4.6433 | 1.1090 |
| RPS21 | P63220 | 4.2661 | 1.4814 |
| SF1 | A0A7P0T9U7 | 4.6589 | 1.0703 |
| MDH1 | A0A5K1VW95 | 4.3828 | 1.3433 |
| RPA3 | P35244 | 4.2243 | 1.4964 |
| DBNL | Q9UJU6 | 4.7122 | 1.0069 |
| NASP | P49321 | 4.1807 | 1.5367 |
| SRSF2 | Q01130 | 3.3334 | 2.3831 |
| PSMB5 | P28074 | 3.5466 | 2.1661 |
| GRN | P28799 | 4.2320 | 1.4765 |
| LYPD3 | O95274 | 4.1342 | 1.5675 |
| EIF3J | O75822 | 4.4644 | 1.2354 |
| RSL24D1 | A0A7I2V3F2 | 4.1154 | 1.5824 |
| ATP5IF1 | Q9UII2 | 4.4906 | 1.2065 |
| CRKL | P46109 | 4.6115 | 1.0738 |
| CRIP1 | P50238 | 3.7836 | 1.9009 |
| PDCD10 | Q9BUL8 | 4.1223 | 1.5456 |
| LYSMD2 | Q8IV50 | 4.0995 | 1.5510 |
| THOP1 | P52888 | 3.7994 | 1.8400 |
| TXNDC17 | Q9BRA2 | 3.2398 | 2.3985 |
| TNKS1BP1 | Q9C0C2 | 4.3840 | 1.2109 |

| | | | |
|---|---|---|---|
| RWDD4 | Q6NW29 | 3.4605 | 2.0924 |
| CARHSP1 | Q9Y2V2 | 4.2285 | 1.3108 |
| TCEAL1 | Q15170 | 3.6791 | 1.8443 |
| PWP1 | Q13610 | 3.8341 | 1.6798 |
| KIAA1143 | Q96AT1 | 3.7246 | 1.7598 |
| UBL5 | Q9BZL1 | 4.0479 | 1.4158 |
| PML | P29590 | 3.8448 | 1.6173 |
| UTP3 | Q9NQZ2 | 4.4393 | 1.0211 |
| PRKCSH | K7ELL7 | 4.1201 | 1.3222 |
| TAGLN2 | P37802 | 3.8682 | 1.5689 |
| GTF2F1 | P35269 | 3.8824 | 1.5438 |
| OR1M1 | Q8NGA1 | 4.1887 | 1.2244 |
| HNRNPD | Q14103 | 3.5170 | 1.8707 |
| ZRANB2 | O95218 | 3.7084 | 1.6783 |
| CKB | P12277 | 4.3035 | 1.0264 |
| CALD1 | Q05682 | 4.1087 | 1.1952 |
| ERP29 | P30040 | 3.9850 | 1.3085 |
| CHLSN | Q9BRJ6 | 3.8825 | 1.4078 |
| LMNB2 | Q03252 | 3.9068 | 1.3833 |
| IGBP1 | P78318 | 4.0867 | 1.1971 |
| COMMD9 | Q9P000 | 4.0156 | 1.2574 |
| CREBBP | Q92793 | 3.6547 | 1.6036 |
| TRAPPC3 | A0A087WWM0 | 3.6837 | 1.5721 |
| CBX5 | P45973 | 3.4458 | 1.8041 |
| YWHAE | P62258 | 2.9396 | 2.3085 |
| HIRIP3 | Q9BW71 | 4.2386 | 1.0058 |
| HDGFL2 | Q7Z4V5 | 4.2068 | 1.0372 |
| CD207 | Q9UJ71 | 3.6824 | 1.5552 |
| SNU13 | P55769 | 3.6313 | 1.6023 |
| PSMD4 | P55036 | 3.2729 | 1.9515 |
| SAFB2 | Q14151 | 3.7774 | 1.4248 |
| MAP1LC3B2 | A6NCE7 | 3.1581 | 2.0410 |
| ATP6V1G1 | O75348 | 3.7578 | 1.4399 |
| DRAP1 | C9JCC6 | 3.9558 | 1.2389 |
| COTL1 | Q14019 | 3.5046 | 1.6887 |
| PDLIM1 | O00151 | 4.1912 | 1.0015 |
| lacZ | P00722 | 3.5131 | 1.6765 |
| HNRNPAB | D6RBZ0 | 3.3329 | 1.8338 |
| KRT19 | P08727 | 3.8955 | 1.2703 |
| PRDX1 | Q06830 | 3.1295 | 2.0233 |
| NUFIP2 | Q7Z417 | 4.0864 | 1.0429 |
| CDKN2C | P42773 | 3.6258 | 1.5035 |
| NHERF2 | Q15599 | 3.7148 | 1.4134 |

| | | | |
|---|---|---|---|
| ADNP | A0A669KBJ7 | 3.7015 | 1.4196 |
| S100A3 | P33764 | 3.2740 | 1.8313 |
| LSM2 | Q9Y333 | 3.4670 | 1.6318 |
| PSME3 | P61289 | 3.7685 | 1.3041 |
| RSRC1 | C9JVB3 | 3.4552 | 1.6130 |
| MAGOHB | Q96A72 | 3.6101 | 1.4536 |
| S100A6 | P06703 | 3.2493 | 1.8098 |
| PSME1 | Q06323 | 3.7173 | 1.3383 |
| ENSA | Q5T5H1 | 3.9242 | 1.1294 |
| RBM3 | P98179 | 3.7866 | 1.2606 |
| PGM1 | A0A3B3ITK7 | 3.3550 | 1.6913 |
| PEPD | A0A494C165 | 3.3193 | 1.7258 |
| CCDC97 | Q96F63 | 3.0648 | 1.9792 |
| RPS28 | P62857 | 3.9146 | 1.1272 |
| UTRN | P46939 | 3.7589 | 1.2731 |
| PIR | O00625 | 3.2609 | 1.7674 |
| SRSF1 | J3KTL2 | 2.9642 | 2.0537 |
| BOLA2B | A0A499FJE1 | 3.2913 | 1.7228 |
| CPSF6 | Q16630 | 3.8916 | 1.1148 |
| TGOLN2 | A0AAG2UW97 | 3.1772 | 1.8235 |
| COPS9 | Q8WXC6 | 3.5156 | 1.4693 |
| ARGLU1 | Q9NWB6 | 3.8230 | 1.1554 |
| YWHAQ | P27348 | 3.2257 | 1.7507 |
| MARCKS | P29966 | 3.5923 | 1.3676 |
| FKBP3 | Q00688 | 3.8682 | 1.0880 |
| TRIR | Q9BQ61 | 2.1060 | 2.8293 |
| S100A16 | Q96FQ6 | 3.4738 | 1.4594 |
| SUMO2 | P61956 | 3.2627 | 1.6665 |
| CAVIN1 | Q6NZI2 | 3.6967 | 1.2307 |
| SF3B5 | Q9BWJ5 | 3.5747 | 1.3393 |
| BRD2 | A0A0G2JK44 | 3.3088 | 1.5795 |
| COMMD1 | Q8N668 | 2.9871 | 1.8966 |
| EWSR1 | Q01844 | 3.5776 | 1.2973 |
| TFG | A0A6Q8PFY7 | 3.1172 | 1.7467 |
| SUB1 | P53999 | 2.9045 | 1.9533 |
| QSOX1 | O00391 | 3.7978 | 1.0345 |
| PLIN3 | O60664 | 3.6883 | 1.1388 |
| SUMO3 | A8MU27 | 3.0946 | 1.7294 |
| SRSF7 | Q16629 | 2.8676 | 1.9395 |
| CWC15 | Q9P013 | 3.6683 | 1.1308 |
| SRP14 | P37108 | 3.4281 | 1.3670 |
| GOLM1 | Q8NBJ4 | 3.5137 | 1.2754 |
| DNAJC9 | Q8WXX5 | 3.6679 | 1.1013 |

| | | | |
|---|---|---|---|
| SCEL | O95171 | 3.7538 | 1.0028 |
| EIF5A | I3L504 | 3.2483 | 1.4900 |
| ANXA8L1 | A0A075B752 | 3.6223 | 1.1152 |
| LRRFIP1 | Q32MZ4 | 3.6792 | 1.0487 |
| MOCS2 | O96033 | 3.4461 | 1.2727 |
| PRDX6 | P30041 | 3.2889 | 1.4102 |
| PIH1D1 | Q9NWS0 | 3.2351 | 1.4432 |
| UBE2N | P61088 | 3.4393 | 1.2378 |
| PEBP1 | P30086 | 2.2975 | 2.3751 |
| UBXN1 | Q04323 | 3.0694 | 1.5901 |
| PPIB | P23284 | 3.2469 | 1.4080 |
| MAP7D1 | Q3KQU3 | 3.2456 | 1.3811 |
| DENR | O43583 | 3.2890 | 1.3282 |
| PPP1R2 | P41236 | 3.1494 | 1.4619 |
| EZR | P15311 | 3.1498 | 1.4498 |
| SRSF3 | P84103 | 2.6888 | 1.9028 |
| TCEA1 | P23193 | 3.3393 | 1.2445 |
| H2A | A0A0U1RRH7 | 3.3182 | 1.2523 |
| ZFAND6 | Q6FIF0 | 3.3487 | 1.2068 |
| HMGN5 | P82970 | 2.8302 | 1.7158 |
| TPI1 | P60174 | 2.4252 | 2.1066 |
| SPC24 | K7ESQ2 | 3.3579 | 1.1596 |
| C1orf198 | Q9H425 | 3.3656 | 1.1505 |
| THOC3 | Q96J01 | 3.0323 | 1.4816 |
| FUS | H3BPE7 | 3.2666 | 1.2457 |
| ITPR1 | Q14643 | 3.3191 | 1.1867 |
| EIF4H | Q15056 | 2.8852 | 1.6108 |
| ZMAT2 | Q96NC0 | 3.2185 | 1.2752 |
| ORC2 | Q13416 | 3.1550 | 1.3344 |
| AK1 | Q5T9B7 | 2.8639 | 1.6064 |
| RBBP4 | Q09028 | 3.3508 | 1.0852 |
| TMSB4X | P62328 | 2.8837 | 1.5384 |
| PSMB3 | P49720 | 2.3675 | 2.0474 |
| EXOSC7 | Q15024 | 3.1788 | 1.2235 |
| ACTN4 | O43707 | 3.2312 | 1.1699 |
| POLR1G | O15446 | 3.3639 | 1.0311 |
| ACYP1 | G3V2U7 | 2.7593 | 1.6240 |
| NCBP2 | P52298 | 3.2859 | 1.0441 |
| GTF2F2 | P13984 | 3.3100 | 1.0166 |
| CLIC1 | O00299 | 2.6995 | 1.6177 |
| CD2BP2 | O95400 | 2.9341 | 1.3820 |
| TP53BP1 | Q12888 | 2.6822 | 1.6327 |
| POLR2J3 | E2QRJ6 | 3.2973 | 1.0106 |

| | | | |
|---|---|---|---|
| CAST | A0A6Q8PH20 | 2.6323 | 1.6714 |
| SDE2 | Q6IQ49 | 3.1812 | 1.1178 |
| ABLIM1 | A0A3B3IS55 | 3.0370 | 1.2353 |
| CREB1 | P16220 | 2.9435 | 1.3267 |
| UBE2I | A0AAA9YHQ8 | 3.2406 | 1.0221 |
| PAFAH1B2 | P68402 | 3.2177 | 1.0396 |
| SFN | P31947 | 2.7032 | 1.5485 |
| RTRAF | Q9Y224 | 2.9611 | 1.2784 |
| STOML2 | Q9UJZ1 | 3.0824 | 1.1535 |
| CPPED1 | Q9BRF8 | 3.1462 | 1.0838 |
| LLPH | Q9BRT6 | 3.0817 | 1.1459 |
| ITPRID2 | P28290 | 3.1636 | 1.0617 |
| PFN1 | P07737 | 2.6637 | 1.5611 |
| NUCB1 | A0A9L9PXH6 | 2.4885 | 1.7133 |
| CALR | P27797 | 2.7256 | 1.4654 |
| KLK10 | O43240 | 3.0422 | 1.1473 |
| GINS4 | Q9BRT9 | 2.9672 | 1.2167 |
| CDC42EP1 | Q00587 | 2.8270 | 1.3412 |
| TXNL1 | O43396 | 2.6862 | 1.4453 |
| PSMB2 | P49721 | 2.0052 | 2.1130 |
| CYCS | P99999 | 2.2459 | 1.8678 |
| CDKN2AIP | Q9NXV6 | 2.9362 | 1.1620 |
| NDUFB10 | O96000 | 2.9938 | 1.0915 |
| PQBP1 | O60828 | 2.7167 | 1.3658 |
| NRDC | O43847 | 2.9967 | 1.0800 |
| PSMB6 | P28072 | 2.4224 | 1.6417 |
| CMPK1 | A0A8V8TMN5 | 2.9368 | 1.1210 |
| LGALS1 | P09382 | 2.7204 | 1.3340 |
| CDK9 | P50750 | 2.8893 | 1.1510 |
| HNRNPA2B1 | P22626 | 2.5445 | 1.4910 |
| H4C1 | P62805 | 3.0315 | 1.0031 |
| PSME2 | A0A087X1Z3 | 2.9666 | 1.0447 |
| ERO1A | Q96HE7 | 2.7833 | 1.2239 |
| CBFB | A0A494C0A9 | 2.7581 | 1.2484 |
| ZNF207 | X6R4W8 | 2.9604 | 1.0418 |
| HNRNPA1 | P09651 | 2.4638 | 1.5358 |
| EIF1AD | Q8N9N8 | 2.6742 | 1.3090 |
| EPS8L1 | Q8TE68 | 2.9382 | 1.0251 |
| P4HB | A0A7P0TA71 | 2.3343 | 1.5988 |
| PPL | O60437 | 2.6403 | 1.2547 |
| REXO4 | Q9GZR2 | 2.8492 | 1.0173 |
| HSPB1 | P04792 | 2.1380 | 1.7123 |
| FAM43A | Q8N2R8 | 2.2979 | 1.5452 |

| | | | |
|---|---|---|---|
| VIM | P08670 | 2.8288 | 1.0125 |
| FDX1 | P10109 | 2.2382 | 1.5855 |
| BAIAP2 | I3L4C2 | 2.7456 | 1.0763 |
| VCP | P55072 | 2.4809 | 1.3151 |
| PCNA | P12004 | 2.5161 | 1.2758 |
| JUNB | P17275 | 2.5839 | 1.1805 |
| DAG1 | Q14118 | 2.5096 | 1.2536 |
| PRDX5 | P30044 | 2.2423 | 1.5181 |
| FHL3 | Q13643 | 2.5608 | 1.1964 |
| IGFBP7 | Q16270 | 2.2615 | 1.4955 |
| CSTF2 | E7EWR4 | 2.6517 | 1.0864 |
| YWHAZ | P63104 | 2.1884 | 1.5461 |
| SDF4 | Q9BRK5 | 2.5606 | 1.1484 |
| YWHAH | Q04917 | 2.5179 | 1.1807 |
| LMNA | A0A6Q8PFJ0 | 2.4853 | 1.2044 |
| MIA3 | Q5JRA6 | 2.5891 | 1.0982 |
| CCDC12 | J3KR35 | 2.6640 | 1.0141 |
| MFAP1 | P55081 | 2.3488 | 1.3059 |
| PITHD1 | Q9GZP4 | 2.5845 | 1.0642 |
| NME1 | P15531 | 2.3561 | 1.2835 |
| RPL23A | A8MUS3 | 2.5294 | 1.1093 |
| MAD2L1 | Q13257 | 2.5857 | 1.0289 |
| CFL2 | Q9Y281 | 2.4964 | 1.1153 |
| PSMA6 | G3V5Z7 | 2.1991 | 1.3795 |
| GCFC2 | P16383 | 2.5383 | 1.0305 |
| B2M | P61769 | 2.2687 | 1.2966 |
| UBE2K | P61086 | 2.3846 | 1.1806 |
| RALB | P11234 | 2.4988 | 1.0652 |
| GLRX5 | Q86SX6 | 2.2109 | 1.3296 |
| SSBP1 | Q04837 | 2.3177 | 1.2029 |
| LGALS3 | P17931 | 2.2459 | 1.2682 |
| KRT17 | Q04695 | 2.2382 | 1.2657 |
| SULT2B1 | O00204 | 2.0929 | 1.3789 |
| YBX1 | P67809 | 2.1531 | 1.3185 |
| ZC3H18 | E7ERS3 | 2.3262 | 1.1267 |
| IST1 | P53990 | 2.2179 | 1.1765 |
| SRFBP1 | Q8NEF9 | 2.3028 | 1.0507 |
| HNRNPH3 | P31942 | 2.3449 | 1.0016 |
| NME1-NME2 | Q32Q12 | 2.2641 | 1.0642 |
| DDB1 | Q16531 | 2.2580 | 1.0309 |
| PPA1 | Q15181 | 2.2296 | 1.0105 |
| PIP | P12273 | 2.0292 | 1.0906 |
| TXNDC5 | Q8NBS9 | 2.0008 | 1.0979 |

| | | | |
|---|---|---|---|
| CFL1 | E9PK25 | 2.0221 | 1.0623 |
